# Physiological fatty acid uptake reveals spatial and systemic constraints on nutrient accessibility in vivo

**DOI:** 10.64898/2026.08.14.744781

**Authors:** Xinyuan Wang, Graham Heieis, Connor Corrigan, Chang Liu, Luuk Reinalda, Laura I. Bogue, Kas Steuten, Kristine Bertheussen, Marouane el Boujadayni, Jeroen M. Punt, Linda V. Sinclair, Mario van der Stelt, Bart Everts, Sander I. van Kasteren, David K. Finlay

## Abstract

Immune cells rely on exogenous fatty acids (FA) for membrane synthesis, bioenergetics and signalling, yet current approaches cannot accurately quantify physiological FA uptake in vivo. Here, we use cyclopropene-tagged fatty acids (cpFA) that, unlike existing FA-uptake tools, are taken up by physiologically relevant mechanisms. We measure FA uptake at single-cell resolution in vivo and uncover a previously unappreciated distinction between nutrient uptake capacity and nutrient accessibility. Although arachidonic acid exhibits the highest uptake capacity ex vivo across immune populations, it displays limited tissue accessibility in vivo, whereas palmitate is broadly accessible. In vivo nutrient-uptake measurements reveal that tissue architecture shapes nutrient accessibility, with spatial constraints in the spleen and exclusion of circulating FA, but not amino acids, from the thymus. Together, these findings identify nutrient accessibility as a distinct layer of metabolic regulation and reveal that immune-cell metabolism is shaped by spatial and systemic constraints on nutrient access

**Highlights:**

- Nutrient accessibility is a distinct layer of metabolic regulation
- Physiological FA uptake differs from ex vivo uptake capacity
- Spatial and systemic factors govern fatty-acid accessibility
- Tissue context shapes immune-cell metabolism in vivo

**In brief:** Using bioorthogonal FA to quantify physiological nutrient uptake in vivo, Wang et al. show that nutrient accessibility is distinct from nutrient uptake capacity. Tissue architecture and systemic FA distribution create spatial constraints on nutrient access, revealing an underappreciated layer of metabolic regulation in immune cells.

## Introduction

Fatty acids (FA) are essential metabolites for immune cells, supporting membrane biogenesis, β-oxidation, lipid-mediator synthesis, and organelle dynamics. The capacity of immune cells to acquire and utilise exogenous FAs is tightly regulated during differentiation, activation, and effector function, and is increasingly recognised as a determinant of immune competence in both physiological and pathological contexts^1,2^. Perturbations in FA metabolism alter T-cell activation, dendritic-cell function, and macrophage polarisation, illustrating the broader principle that metabolic programming is integral to immune fate and effector potential^3^. Despite this central role, methods for quantifying FA uptake at single-cell resolution remain limited. Common fluorescent FA analogues, such as BODIPY-conjugated long-chain FAs, are dominated by the hydrophobic dye rather than the FA moiety, leading to uptake patterns that do not reflect physiological transport^4^. As a result, fundamental questions about FA acquisition across immune lineages, activation states, and tissue microenvironments remain unresolved.

A growing body of work highlights that immune-cell metabolism measured *ex vivo* or *in vitro* often diverges markedly from metabolic behaviour *in vivo*^5–7^. Tissue microenvironments impose constraints in oxygen tension, nutrient distribution, stromal interactions, and metabolite gradients that cannot be recapitulated in standard culture media. Immune cells in inflamed tissues or tumours compete with stromal and parenchymal cells for glucose, amino acids, and lipids, creating metabolic pressures invisible to isolated-cell assays ^8^. Moreover, nutrient and oxygen gradients vary sharply across short spatial scales in tissues^9^, causing cells of identical lineage to adopt distinct metabolic programs depending on their microenvironment ^10,11^. During immune responses and disease progression, these constraints shift dynamically. Thus, metabolic phenotypes measured after cell isolation can misrepresent the true metabolic state of immune cells in situ, obscuring the extent to which nutrient accessibility is shaped by tissue architecture, vascular supply and local competition. This highlights the need for technologies capable of directly measuring nutrient acquisition in vivo and at single-cell resolution.

Bioorthogonal click chemistry provides a powerful strategy for labelling biomolecules in situ without interfering with endogenous biochemical pathways and has become widely used for cell-surface and metabolic tracing applications^12^. The key conceptual advance of this approach has been the use of small, inert handles that allow fluorophores to be attached after uptake, eliminating the artefacts inherent to pre-labelled nutrient probes^13^.

Building on these principles, we recently developed cyclopropene-tagged FA (cpFAs)^14,15^. These are analogues of FA in which a cyclopropene has been introduced. By placing it across a double bond, these FAs are only one carbon different from the unmodified nutrients. In addition, the cyclopropene -group is a bioorthogonal group, which means it can be selectively reacted with electron-poor diene in biological medium without the need for additional catalyst^16^. This allows the fluorescent quantification of the cyclopropene-modified FA by uncoupling FA transport from fluorophore detection; cpFAs enable measurement of FA acquisition with minimal perturbation of endogenous transport and metabolic handling.

A major gap in the field is the lack of tools that can distinguish between what immune cells can take up from what FA they can access in vivo. Most uptake assays are performed ex vivo, where isolated cells are given equal access to nutrients. This does not capture the effects of tissue architecture, blood supply, anatomical barriers, or competition for nutrients that is encountered in vivo. As a result, even fundamental questions about in vivo FA uptake remain unanswered, including which immune cell populations acquire circulating FA and how tissue-specific metabolic niches shape FA availability and preference. Methods capable of directly measuring FA acquisition in vivo at single-cell resolution are therefore needed.

Here, we introduce and apply a panel of cpFA probes to interrogate FA acquisition across immune lineages ex vivo and in vivo. Using this platform, we identify intracellular pathways that support FA uptake in vivo, and we resolve tissue- and lineage-specific preferences in FA acquisition, hereby uncovering previously inaccessible features of nutrient accessibility, including the influence of tissue architecture, vascular organization and anatomical barriers. Together, these findings reveal how intrinsic metabolic capacity and local tissue environments interact to shape immune-cell access to FA in vivo.

## Results

### Dynamic cpOA uptake with immune development and activation

We previously synthesized a panel of cpFAs and demonstrated that they can enter immune cells in vitro^14^. Here, we sought to determine whether these cpFAs are taken up through physiological FA transport pathways, rather than via non-specific membrane insertion or other artefactual routes. We focused on cyclopropene-oleic acid (cpOA) as a representative probe. To establish temperature dependence, a key indicator of protein-mediated transport, cpOA uptake was quantified in human NK92 (NK cell), THP1 (monocyte/macrophage), and HEK293T (epithelial) cell lines at 37 °C versus 4 °C. All three lines showed significantly reduced uptake at 4 °C, consistent with transport protein-dependent FA import (Fig. 1A,B). After establishing the dose-signal ratio, kinetics of uptake, and a concomitant optimal concentration range and uptake interval for subsequent studies (Fig. 1C,D; Supplementary Fig. 1A,B), we next measured cpOA uptake capacity in primary murine immune cells using ex vivo splenocytes. This uptake capacity varied substantially across lymphoid and myeloid subsets, with highest incorporation in macrophages and dendritic cells (Fig. 1E-G). Because metabolic activation increases FA demand, we also examined changes in cpOA uptake capacity upon immune cells activation. For this, murine NK cells were activated *in vivo* following intraperitoneal injection of Poly(I:C) (10 mg/Kg) and examined ex vivo 18 hours after this challenge. Activated NK cells exhibited significantly increased cpOA uptake capacity compared to resting cells (PBS injection control) (Fig. 1H), suggesting that circulating FA are a fuel source to support the increased cellular growth and energy consumption in these activated innate lymphocytes.

**Figure 1.**
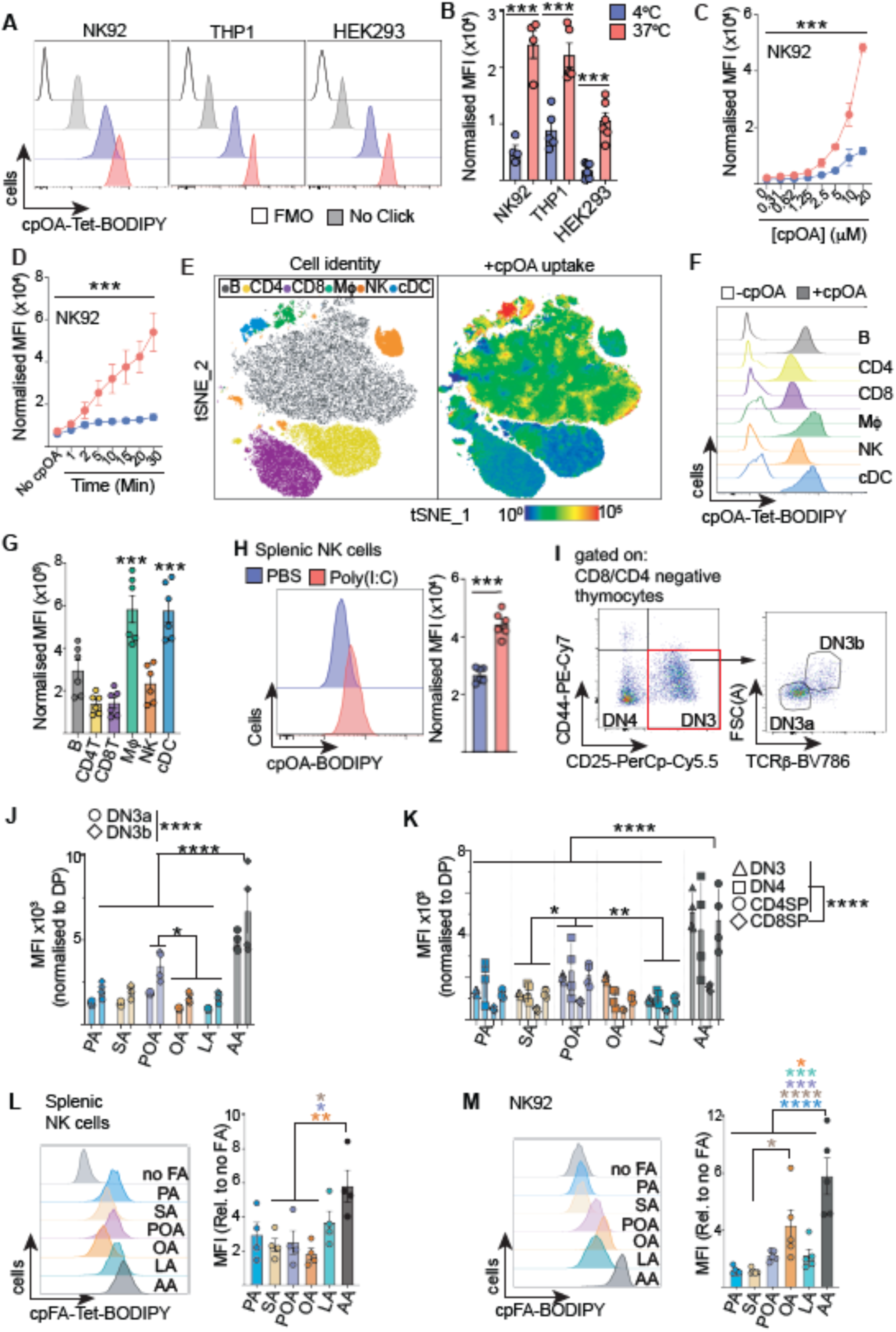
Uptake of cpFA probes is quantified across immune development and activation. **(A,B)** NK92-MI, THP-1, and HEK293T cells were incubated with cpOA (10 μM) for 10 min at 37°C or 4°C, followed by fixation and Tetrazine-BDP click reaction. FMO and no cpOA controls are shown (A). Pooled cpOA uptake data presented MFI, adjusted to subtract no-cpOA background fluorescence. **(C,D)** cpOA uptake assays were performed in NK92 cells for 10 min using increasing concentrations of cpOA, as indicated (C), and using (10 μM) cpOA and increasing incubation times, as indicated (D). **(E)** t-distributed stochastic neighbor embedding (t-SNE) analysis of splenocytes following 10 min cpOA uptake (10 μM). Left, annotated splenocyte populations; CD19+ B cells, CD3+CD4+ T cells, CD3+CD8+ T cells, CD3-CD19-F4/80+CD11c+ macrophages, CD3-CD19-NK1.1+ NK cells, and CD3-CD19-CD11c+MHCII+ cDC. Right, corresponding cpOA uptake intensity. **(F,G)** cpOA uptake analysis, cpOA (10 μM) for 10 min, using splenocytes and antibody staining to identify immune subsets. Representative histograms showing cpOA uptake and corresponding no-cpOA background controls (F). Pooled data showing cpOA uptake MFI for immune subsets, adjusted for no-cpOA background. (**H)** Mice received i.p. Poly(I:C) (10 mg/kg) and splenic (CD3-CD19-NK1.1+) NK cells analysed after 18 hours for cpOA uptake. Representative histogram (left) and pooled data of background adjusted MFI (right). **(I-K)** Thymocytes were isolated and uptake of 6 different cpFAs (10 μM for 10 min) measured across β-selection checkpoint, DN3a to DN3b subsets (I). Pooled data showing uptake of 6 cpFA adjusted for background and normalised to CD4 CD8 double positive thymocytes (J). Uptake of 6 cpFAs measured for DN3, DN4, CD4+ single positive (SP) and CD8+ SP thymocyte subsets, normalised to DP thymocytes (K). (**L,M,N)** Uptake of 6 cpFA (10 μM for 10 min) into splenic NK cells (L) and NK92 cells (M) adjusted for background, showing representative histograms (L,M, left) and pooled data (L,M, right). Data is representative (A,E,F,H,I, L,M) or mean +/- SEM (B-C,G,H,J,K,L,M) for 4-6 independent experiments (A-D,M); for 4 (I-L) o 6 (E-H) mice across a least 3 independent experiments . Statistical analysis was performed using ANOVA or non-parametric tests as appropriate, with Šidák (B,J,K) or Tukey (G,L,M) post-tests. ns, not significant; *p < 0.05; **p < 0.01; ***p < 0.001; ****p < 0.0001; SA, steric acid; PA, palmitic acid; OA, oleic acid; POA, palmitoleic acid; LA, linoleic acid; AA, arachidonic acid.

The metabolic orchestration of thymocyte development remains a major unresolved challenge; nutrient dependencies of the dynamic developmental stages that underpin T-cell maturation have not been fully resolved. Notably, endogenous oleic acid biosynthesis has recently been implicated in thymocyte maturation^17^. Therefore, we investigated the uptake of our recently described panel of 6 cpFA in thymocyte subsets. This panel includes cyclopropene-containing analogues of saturated FA (SFA), steric acid (cpSA), palmitic acid (cpPA); monounsaturated FA (MUFA), palmitoleic acid (cpPOA) and cpOA, as well as polyunsaturated FA (PUFA) linoleic acid (cpLA) and arachidonic acid (cpAA) (Supplementary Fig. 1C)^14^. Initially, the uptake of these 6 different cpFA were measured at a critical checkpoint in thymocyte development called β-selection. β-selection occurs when thymocytes successfully rearrange their T cell receptor (TCR) β chain locus. Expression of TCRβ along with the pre-Tα chain and other signalling components, allows for functional pre-TCR signalling, which drives a metabolic burst, robust proliferation and differentiation from small TCRβ negative DN3a to large TCRβ positive DN3b (Fig. 1I). DN3a to DN3b differentiation was associated with significant increase in cpFA uptake, in particular cpAA and to a lesser extent cpPOA (Fig. 1J). Higher uptake of cpAA relative to other cpFA was observed across all thymocyte subsets (Fig. 1K). Other differences measured including lower cpFA uptake into CD8+ single positive (SP) thymocytes compared to DN and CD4 SP thymocytes (Fig. 1K). The uptake of these 6 cpFA was also measured in splenic NK cells (Fig. 1L,M) and in human NK92 cells (Supplementary Fig.1E), and both showed a greater preference for uptake of cpAA compared to uptake of the other cpFA.

### cpOA uptake reflects physiological FA transport

Boron dipyrromethene (BODIPY)-C16 conjugates are fluorescently conjugated C16 FA widely used to assess FA uptake in immune cells ^18^. However, because the bulky, highly hydrophobic BODIPY moiety can dominate the molecule’s biophysical properties, its cellular accumulation may not reflect bona fide FA transport. To directly compare cpOA and BODIPY-C16 uptake, NK92 cells were incubated at either 37 °C or 4 °C and their intracellular fluorescence distribution was assessed. Representative images are shown in Fig. 2A, and pooled single-cell fluorescence measurements are shown in Fig. 2B. To estimate fluorescence associated with active FA transport, values were normalized to the corresponding 4 °C controls. cpOA uptake was significantly reduced at 4 °C, whereas the stronger BODIPY-C16 signal was unaffected by temperature. Because the intense BODIPY-C16 fluorescence obscured the temperature-dependent differences in cpOA signal in Fig. 2A, additional images were acquired without BODIPY-C16 and with longer exposure times (Fig. 2C). These images clearly show temperature-dependent cpOA uptake and include no-cpOA and no-fluorophore controls. These observations were corroborated by flow cytometry, which showed that cpOA uptake was highly temperature-dependent, whereas BODIPY-C16 generated strong fluorescence signals that were largely unaffected by temperature (Fig. 2D,E). Because both transporter-mediated FA uptake and intracellular metabolic trapping are strongly suppressed at 4 °C, physiological FA uptake is expected to be minimal under these conditions. The maintenance of high BODIPY-C16 fluorescence at 4 °C therefore indicates that a substantial proportion of the signal does not arise from physiological FA uptake. Rather, BODIPY-C16 appears to accumulate through temperature-insensitive mechanisms that are unlikely to reflect endogenous FA transport. These findings highlight an important limitation of BODIPY-conjugated FA probes for measuring FA uptake in immune cells.

**Figure 2.**
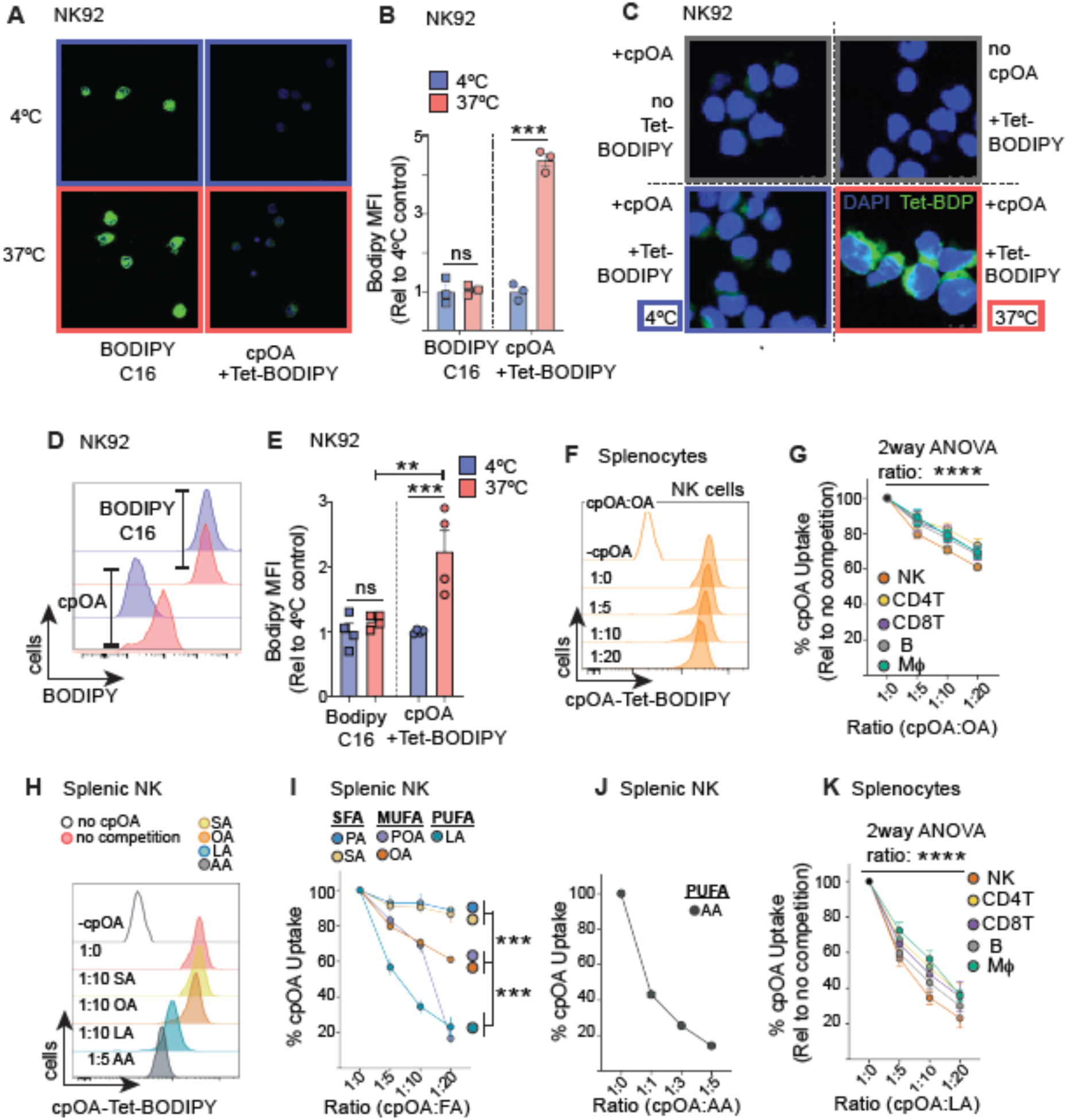
cpOA uptake, but not that of C16-BODIPY, uses physiological pathways. **(A-E)** NK92 cells were treated with cpOA (10 μM) or BODIPY-C16 (10 μM) for 10 min at 37°C or 4°C. Tet-BODIPY was attached to cpOA after cell fixation and permeabilization and cells analysed by confocal microscopy (A-C) and flow cytometry (E,F). Representative images showing BODIPY fluorescence (green) and nuclei stained with DAGP (blue) **(A)** and pooled quantitation of fluorescence at 37°C relative to 4°C control **(B).** Representative images showing cpOA uptake at 37°C or 4°C with Tet-BODIPY conjugation and additional no cpOA and no Tet-BODIPY controls **(C).** Representative histogram of cpOA+Tet-BODIPY versus C16-BODIPY fluorescence after 37°C or 4°C uptake incubations **(D)** and associated pooled data for MFI, shown relative to 4°C controls **(E). (F-K)** Splenocytes were incubated with cpOA (10 μM) for 10 min at 37°C in the presence of varying concentrations of native OA or native forms of other FA. Representative histogram of cpOA uptake into splenic NK cells in the presence of increasing molecular ratios of native OA from 1:0 (no native OA) to 1:20 (20-fold excess native OA) **(F).** Pooled data for cpOA uptake into splenic subsets as labelled and increasing ratios of native OA, shown relative to 1:0 ratio **(G).** Representative histograms of cpOA uptake into splenic NK cells with different FA species as labelled **(H)** and pooled data showing competition of cpOA uptake by increasing ratios of different FA species **(I)** with AA shown separately as potent competition required different ratios **(J).** Pooled data showing competition for cpOA uptake by LA at increasing ratios for different splenic subsets, shown relative to 1:0 **(K).** Data is representative (A,C,D,F,H) or mean +/- SEM (B,E,I-K) for 3 independent experiments and 1×10^6^ cells per experiment (B), 4 independent experiments (D,E) and for 4-6 spleens across 4 independent experiments (F-K). Statistical analysis was performed using ANOVA or non-parametric tests as appropriate; Šidák (B,E) or Tukey (I,J) post-tests. ns, not significant; **, p < 0.01; ***, p < 0.001; ****, p < 0.0001; SA, steric acid; PA, palmitic acid; OA, oleic acid; POA, palmitoleic acid; LA, linoleic acid; AA, arachidonic acid.

To verify that cpFA uptake occurs via physiologically relevant routes, we performed competition experiments with unlabeled FA. We tested if the cpOA uptake signal was reduced when an excess of unmodified OA was present. In murine splenic NK cells, there was a dose dependent decrease in cpOA associated fluorescence when unmodified OA was included up to a 20:1 excess (Fig. 2F,G). Similar decreases were observed for other splenic subsets including T cells, B cells and macrophages (Fig. 2G); at a 1:20 ratio of cpOA:OA there was a decrease of 25-40% of cpOA uptake, which was less than the expected complete block at this excess of unmodified FA. The affinity of FA species for binding to proteins linked to FA uptake varies with strongest binding described for certain PUFAs^19,20^. We therefore considered if PUFA might be superior competitors for cpOA uptake, thus providing further evidence of physiological cpOA uptake. Indeed, the PUFA linoleic acid (LA) was a potent competitor, reducing cpOA uptake into splenic NK cells by up to ∼80% with a 1:20 molar excess of cpOA:LA (Fig. 2H,I). MUFAs OA and palmitoleic (POA) showed moderate inhibition of cpOA uptake (∼30% reduction), saturated FA (SFAs) palmitic and stearic acid (PA and SA) showed minimal competition (Fig. 2H,I, Supplementary Fig. 2A). The PUFA arachidonic acid (AA) was striking in its ability to compete for cpOA uptake, achieving >80% inhibition when only at 5-fold molar excess (1:5; cpOA:AA)(Fig. 2H,J, Supplementary Fig. 2B). Beyond splenic NK cells, similar patterns were observed across other splenic subsets; shown is LA competition for cpOA for T cells, B cells and macrophages (Fig, 2K).

These results demonstrate that cpOA uptake is a regulated process that is temperature-sensitive and competed by physiological unmodified FA, consistent with engagement of endogenous FA transport machinery. This contrasts with the non-physiological accumulation of BODIPY-C16. Thus, cpFA probes provide a more accurate means of quantifying physiologically relevant FA uptake in immune cells.

### FABP5 is key to facilitating cpFA uptake into immune cells

To identify factors that support cpFA uptake in immune cells, we correlated cpOA uptake across immune subsets with known abundances of known FA-handling proteins. Protein copy numbers were obtained from public datasets (immpres.co.uk) and in-house proteomic data^21,22^. We focused on FA transport proteins (SLC27/FATP family), FA binding proteins (FABP family), and scavenging receptors such as CD36 (Fig. 3A). Among all candidates, FABP5 showed the strongest correlation with cpOA uptake (Pearson r=0.81, p = 0.001), with other significant associations for SLC27A1 and FABP4 (Supplementary Fig. 3A)(Fig. 3B). While CD36 expression did not correlate with observed cpOA uptake signals it should be remembered that CD36 is not uniformly expressed for many immune subsets (Fig.3B). To test whether FABP5 is required for cpFA uptake, we disrupted FABP5 expression in NK92 and THP-1 cells using CRISPR-mediated insertion of a selection cassette into exon 1 (Supplementary Fig. 3B)^23^. FABP5-knockout (FABP5^KO^) clones were isolated, expanded, and validated by flow cytometry and Western blotting (Fig. 3C; Supplementary Fig. 3C). All FABP5^KO^ THP-1 clones exhibited a consistent and reproducible reduction in cpOA uptake compared with wild-type (FABP5^WT^) cells (Fig. 3D). Similar results we obtained in FABP5^KO^ NK92 cells (Supplementary Fig. 3D,E). When the full panel of cpFAs was tested, loss of FABP5 expression affected the uptake of those cpFA with the highest uptake signals, including cpOA, cpLA and cpAA (Supplementary Fig. 3F-H).

**Figure 3:**
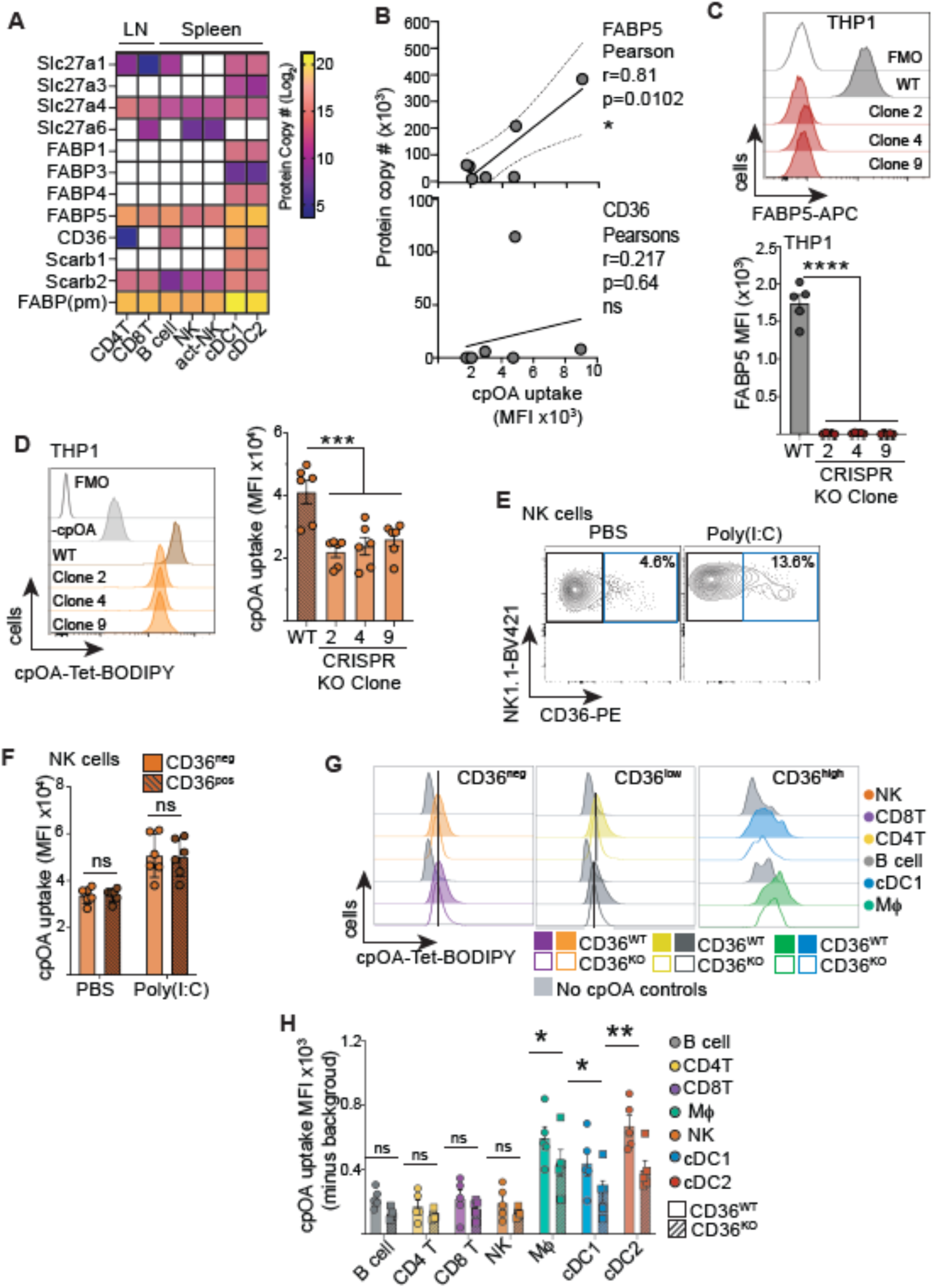
FABP5 and CD36 contribute to cell type-specific cpOA uptake. **(A,B)** Protein copy numbers for proteins associated with FA transport for murine splenocyte populations and lymph node T cells from proteomic datasets (immpres.co.uk, PRIDE), presented on a heatmap showing low (blue) to high (yellow) expression (A) and correlated for CD36 and FABP5 with corresponding cpOA uptake measured by flow cytometry (B). **(C,D)** Representative histogram and pooled data showing FAPB5 expression (C) and cpOA uptake (D) in WT THP-1 and 3 independent clones of FABP5-/- THP-1 single-cell clones. **(E,F)** Mice were injected i.p. with Poly(I:C) (10 mg/kg) and splenic NK cells analysed for CD36 expression after 18 hours (E) and the uptake of cpOA in CD36 negative versus positive NK cells measured (F). **(G,H)** cpOA uptake into splenocytes from CD36 WT and CD36 KO mice was measured and representative histograms shown (G) and pooled data (H). Data is representative (C,D,E,G), mean (A,B), mean +/- SEM (C,D,F,H) for 3-4 proteomic and uptake replicates (A,B), 5-6 independent experiments (C,D) and 4 independent experiments (G,H). Statistical analysis was performed using ANOVA or non-parametric tests as appropriate; Šidák (H,F) or Dunnett’s (C,D) post-tests or using a Pearson’s test (B). ns, not significant; *, p < 0.05; **, p < 0.01; ***, p < 0.001; ****, p < 0.0001.

Because CD36 has been widely implicated in FA uptake in multiple immunological contexts, we next examined its contribution to cpOA uptake^24,25^. CD36 is minimally expressed on most lymphocyte subsets in the steady state, but a small proportion of NK cells upregulate CD36 following *in vivo* activation (Fig. 3E). In these splenic NK cells, cpOA uptake did not differ between CD36⁺ and CD36⁻ NK cells, indicating CD36-independent cpOA uptake in these lymphocytes. In contrast, some myeloid cells including conventional dendritic cells (cDC) and macrophages express high levels of CD36 at rest and are among the immune subsets with the highest cpOA uptake (Fig. 1C). To directly test CD36 function, we compared cpOA uptake in splenocytes from CD36^WT^ and CD36^KO^ mice. cpOA uptake was unchanged in B cells, T cells and NK cells from CD36^KO^ mice. However, cDC1, cDC2, and macrophage subsets from CD36^KO^ mice showed significantly reduced cpOA uptake (Fig. 3H). This demonstrates that CD36 contributes to cpFA uptake in certain CD36-expressing myeloid, but not lymphoid populations.

Together, these data identify FABP5 as an important intracellular FA-handling protein in immune cells required for efficient cpFA uptake, while CD36 supports cpFA acquisition in specific myeloid subsets.

### cpFA uptake by CD4⁺ Tregs is both tissue- and lineage-dependent

Regulatory T cells (Tregs) are essential for maintaining immune homeostasis and, *in vitro*, display increased reliance on fatty-acid oxidation compared with naïve or effector T (Teff) cells ^26^. However, whether Tregs preferentially acquire specific FA (FAs) in different *in vivo* niches remains unclear, despite recent evidence implicating oleic acid in thymic Treg development ^17^. To address this, we profiled uptake of representative saturated (cpPA), monounsaturated (cpOA), and polyunsaturated (cpAA) FA across CD4⁺ T-cell subsets isolated from multiple tissues. To enable direct comparison between tissues, CD3⁺ T cells were enriched *ex vivo* and barcoded using tissue-specific anti-CD45 fluorophore conjugates prior to pooling and incubation with cpFAs. This approach ensured that all CD4⁺ T cells experienced identical culture conditions, minimizing technical and tissue-specific bias (Fig. 4A). Using this strategy, we assessed FA uptake differences (i) between tissues, (ii) between CD4⁺ T-cell subsets within each tissue, and (iii) across FA classes (SFA, MUFA, PUFA).

**Figure 4.**
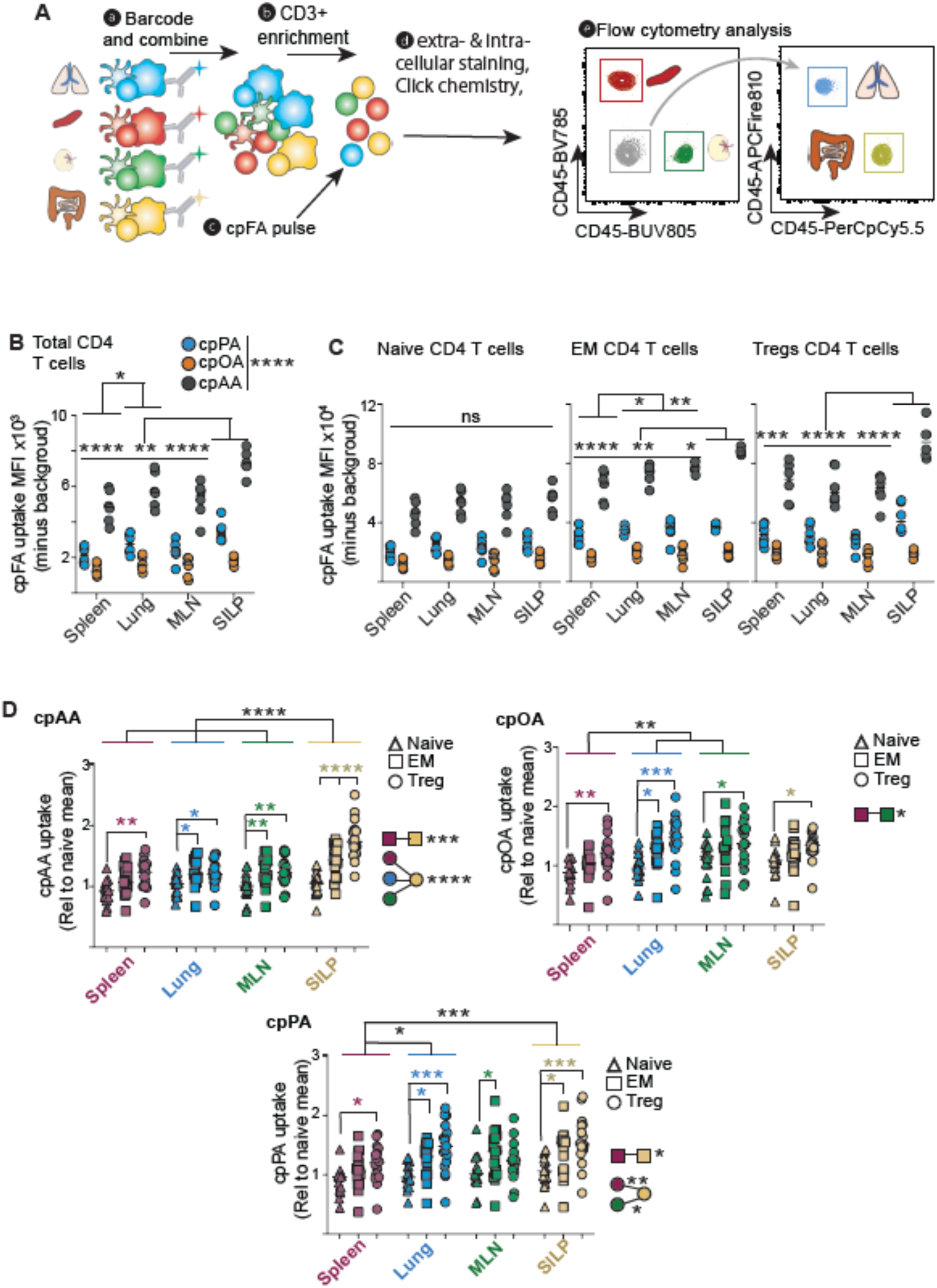
Tissue residency shapes FA acquisition by regulatory T cells. **(A-D)** T cells were enriched from spleen, MLN, ALN and small intestine lamina propria using CD3+magnetic bead sorting, and each stain with anti-CD45 tagged with a distinct fluorophore, then combined. Combined CD4 T cells were incubated with 25 μM cpOA, cpPA, cpAA or PBS for 30 min at 37°C, cells prepared for flow cytometry analysis (A). Uptake of each cpFA into total CD4 T cells was compared for each tissue location (B). CD4 T cells were stratified by cell type into naïve, effector memory and Treg CD4 T cells and uptake of different cpFA analysed (C). Data was separated into the different cpFA species and compared across tissues and CD4 T cells subsets (D). Data shows individual uptake experiments + median for 6 mice representative of 3 individual experiments (B,C), 17 mice over 3 individual experiments (D). Statistical analysis was performed using ANOVA and Tukey post-tests. ns, not significant; *, p < 0.05; **, p < 0.01; ***, p < 0.001; ****, p < 0.0001.

At the global level, total CD4⁺ T cells from the small-intestine lamina propria (SILP) exhibited the highest cpFA uptake (cpAA and cpPA), followed by CD4⁺ T cells isolated from mesenteric lymph nodes (MLN) or spleen (Fig. 4B). Uptake of the 3 cpFAs into total CD4+ T cells was significantly between tissues (Fig.4B). This could arise either because tissues contain different proportions of naïve, EM, and Treg cells, or because the same CD4⁺ T-cell subset exhibits different FA uptake capacities depending on its tissue of residence. To distinguish between these possibilities, total CD4⁺ T cells were subdivided into naïve (CD44⁻), effector/memory (EM; CD44⁺), and regulatory (Foxp3⁺) populations. Naïve CD4⁺ T cells displayed comparable uptake of cpPA, cpOA, and cpAA across all tissues examined (Fig. 4C). In contrast, both EM and Treg populations exhibited pronounced tissue-specific differences. EM and Treg CD4⁺ T cells from the SILP showed greater uptake of all cpFAs compared with corresponding EM and Treg populations from spleen, lung, and MLN, while lung and MLN EM cells also demonstrated higher uptake than splenic EM cells (Fig. 4C).

To further interrogate FA-specific uptake patterns, pooled data from three independent experiments were normalized to total naïve CD4⁺ T-cell MFI and analyzed by FA class (Fig. 4D). Uptake of the PUFA cpAA was highest overall (Fig. 4B,C), with SILP CD4⁺ Treg cells exhibiting the highest cpAA uptake (Fig. 4D, left top panel). EM CD4⁺ T cells also had elevated cpAA uptake relative to naïve cells in the SILP, MLN, and lung, albeit lower than the Treg cells (Fig. 4D, left top panel). In contrast, uptake patterns for the MUFA cpOA differed from PUFA cpAA uptake patterns (Fig. 4D, top right panel). Tregs consistently transported more cpOA than naïve CD4⁺ T cells within each tissue, cpOA uptake was greatest in Tregs from lung and MLN. CD4^+^ EM T cells from lung and MLN also showed increased uptake of cpOA relative to naïve cells. Finally, we looked at the uptake pattern of the SFA, cpPA. The Tregs showed greatest cpPA uptake compared with naïve CD4^+^ T cells, with the biggest differences seen in SILP, spleen, and lung (Fig. 4D, bottom panel). CD4^+^ EM T cells also showed enhanced cpPA uptake over naïve counterparts across all tissues. Notably, both EM and Treg populations from the SILP demonstrated higher cpPA uptake capacity compared with their splenic counterparts.

Collectively, these data reveal pronounced tissue- and subset-specific differences in cpFA uptake, with additional discrimination across FA classes. Overall, CD4⁺ T cells exhibited a preferential uptake hierarchy of cpAA > cpPA > cpOA, a pattern most prominent within the SILP, where both EM and Treg populations displayed enhanced capacity for PUFA and SFA uptake.

### In vivo uptake of cpOA reveals organ-specific access to circulating FA

*Ex vivo* profiling experiments demonstrated the utility of the cpFA probe library for dissecting immune cell FA uptake capacity (Fig. 3,4); however, *in vivo* measurements are required to capture additional layers of complexity imposed by tissue architecture, blood flow, nutrient gradients, and tissue-specific metabolic niches. To assess *in vivo* applicability, we administered cpOA intravenously and quantified uptake in circulating and splenic immune cells relative to PBS-injected controls (Fig. 5A,B).

**Figure 5.**
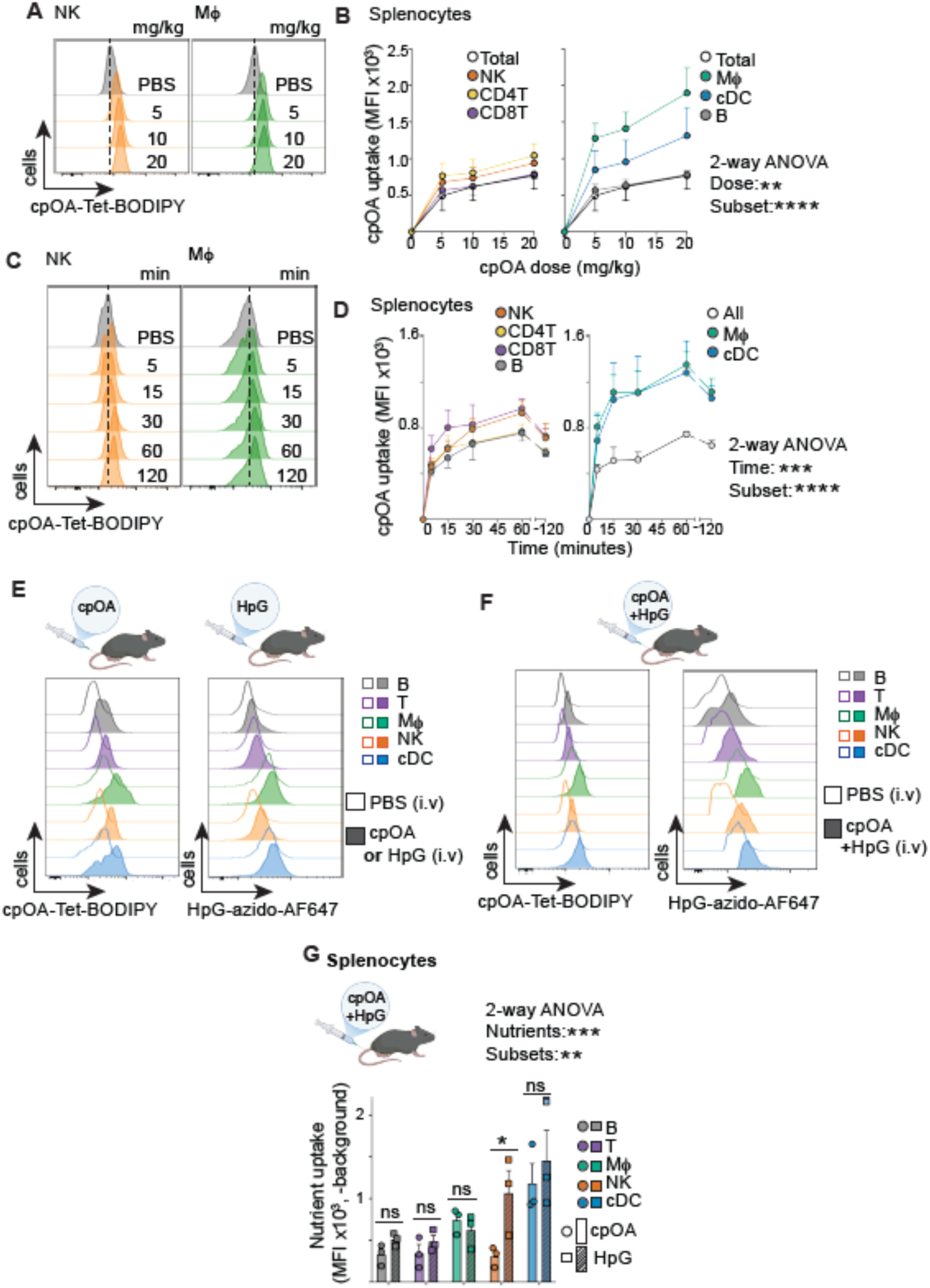
In vivo characterization of cpOA uptake across immune cell populations. **(A-D)** Mice were i.v. injected with cpOA at indicated doses (A,B) and with 10 mg/kg for indicated times (C,D) or PBS controls, and spleens isolated, cells prepared for flow cytometry analysis. Representative histograms are shown for NK cells and macrophages (Mϕ)(A,C) and pooled data for total splenocytes; T cells and NK cells (left); and B cells, Mϕ, and cDC (right) (B,D). (**E-H)** Mice were i.v. injected with 10 mg/kg cpOA, 100 mg/kg HPG,10 mg/kg cpOA and 100 mg/kg HPG together, or PBS control; then after 30 min spleen and thymus were isolated and cells prepared for flow cytometry analysis. **(E,F)** Representative histograms showing *in vivo* uptake of cpOA (left) or HPG (right) in separate mice (E) or the same mice after co-injection showing different splenocyte populations. **(G)** Pooled data showing in vivo uptake of cpOA and HpG in splenic populations. Data is representative (A,C,E,F)) and mean +/- SEM (B,D,G) for 3 (A-G) and 6 (H) individual experiments. Statistical analysis was performed using ANOVA or non-parametric tests as appropriate; Šidák post-tests (G). ns, not significant; *, p < 0.05; **, p < 0.01; ***, p < 0.001; ****, p < 0.0001.

Dose titration experiments identified that the lowest dose 5 mg/kg of cpOA was reliably detectable *in vivo*, and 10 mg/kg as the optimal dose that provided robust signal without excessive injection viscosity (Fig. 5A,B; Supplementary Fig. 4A,B). Assuming about 2 ml blood volume per mouse the concentration of cpOA in blood after injection was ∼350 μM, which is below the report concentration for OA in mouse serum^27^. At this dose, cpOA uptake in blood and splenic lymphocytes, as well as splenic myeloid cells, was detectable within 5 minutes post-injection and reached an apparent plateau by 30 minutes (Fig. 5C,D; Supplementary Fig. 4C,D). Time-dependent uptake in blood NK and B cells was well described by a hyperbolic function (R^2^ = 0.973 and 0.984, respectively) and was consistent across biological replicates (Supplementary Fig. 4E). This kinetic profile indicates rapid initial uptake followed by a progressive reduction in net accumulation rate, consistent with a finite cellular uptake capacity and/or limited nutrient availability.

We next asked whether the in vivo uptake of FA and amino acids could be measured simultaneously. To achieve this, we combined cpOA with L-homopropargylglycine (HpG), a clickable amino acid probe that reports uptake through the glutamine transporter SLC1A5^13^. The distinct chemical handles present on cpOA and HpG permit selective labeling with different click reactions and fluorophores, enabling simultaneous quantification of FA and amino acid uptake in individual cells. Mice received cpOA alone, HpG alone, or both probes together, and nutrient uptake was analyzed in splenic immune cells 30 min later. Comparable uptake signals were obtained whether each probe was administered alone or in combination (Fig. 5E-G, Supplementary Fig. 4F,G), validating this multiplexed nutrient-tracing approach. Across splenic immune-cell subsets, incorporation of HpG and cpOA was broadly similar, with the notable exception of NK cells, which showed preferential acquisition of glutamine (HpG) over FA (cpOA) (Fig. 5G). Importantly, this pattern was observed both when the probes were administered separately and when they were co-injected into the same mouse (Fig. 5E-G), indicating that simultaneous nutrient tracing and sequential click labeling do not materially affect quantification of either probe.

### Spatial factors influence *in vivo* cpOA uptake in the spleen

We next investigated whether tissue architecture and spatial positioning influence cpOA acquisition by immune cells *in vivo*, and contrasted this with uptake of HpG, a clickable glutamine analogue transported into cells via the amino-acid transporter Slc1a5 ^13^. A key distinction between these probes lies in their mode of delivery: HpG is a small, freely diffusible amino acid, whereas cpOA is a substantially larger hydrophobic FA molecule that circulates bound to the 67kDa protein albumin. Consequently, cpOA and HpG are expected to differ fundamentally in how readily they access immune cells within intact tissue. To test this, we compared cpOA and HpG uptake patterns measured in intact spleens following in vivo delivery with uptake measured ex vivo immediately after mechanical dissociation. In this situation, spatial constraints are removed and all cell types have equal access to substrate. As expected, uptake measurements of both probes were lower in vivo than ex vivo (Fig. 6A,B). This difference highlights an important conceptual distinction: ex vivo assays measure the capacity of cells to acquire nutrients when access is unrestricted, whereas in vivo measurements capture physiological nutrient acquisition, which is shaped by nutrient availability, vascular delivery, tissue architecture, local competition and endogenous nutrient pools (Fig. 6A,B).

**Figure 6:**
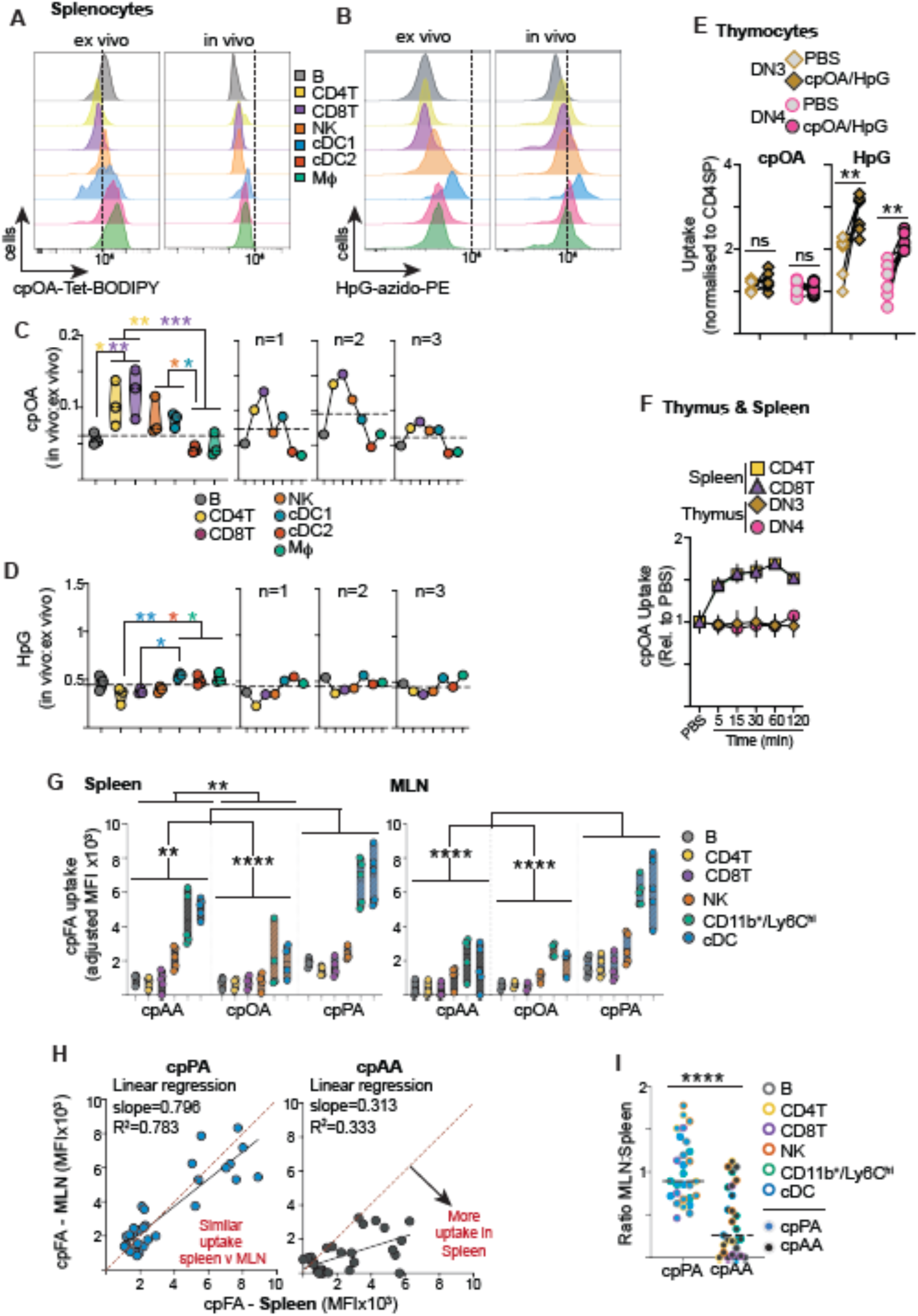
Tissue architecture and systemic FA distribution shape nutrient accessibility in vivo. **(A-F)** Mice were i.v. injected with 10 mg/kg cpOA (A-F), 100 mg/kg HPG (A-E),1 or PBS control. Spleens and thymuses were isolated, cells prepared for flow cytometry analysis. Splenocytes from PBS injected mice were prepared as in vivo uptake controls and used for ex vivo cpOA and HpG ex vivo uptake assays. (**A,B**) Representative histograms of in vivo and ex vivo uptake for splenocyte subsets for cpOA (A) and HpG (B) uptake. Dotted lines indicate MFI values at MFI = 10^4^. **(C,D)** Ratio of in vivo:ex vivo uptake MFIs were calculated and pooled data (left) and ratios for each experimental repeat (right) are shown for cpOA (C) and HpG (D) uptake. Dotted lines indicate the mean of ratios across all subsets for pooled (left) and individual experiment (right). (**E**) Pooled data showing in vivo uptake of cpOA (left) and HpG (right) in DN3 (diamond) and DN4 (circle) thymocytes compared to PBS controls. **(F)** Splenocytes and thymocytes were isolated after increase time periods after i.v injection of 10 mg/kg cpOA (5 min to 120 min) and in vivo cpOA uptake measured in CD4 (square) and CD8 (triangle) splenic T cells and DN3 (diamond) and DN4 (circle) thymocytes relative to control mice injected with PBS. **(G-J)** Mice were i.v. injected with 34 μmoles of cpOA, cpPA, or cpAA and spleen and MLN harvested after 60 min. Cells were isolated and prepared for flow cytometry analysis (E). Pooled data for in vivo uptake of each cpFA into T cells, B cells, NK cells, cDC and CD11b^+^/Ly6C^high^ monocytes isolated from the spleen or MLN, shown as background (PBS) adjusted MFI values (F). cpFA uptake (adjusted for PBS background) for cpAA and cpPA was plotted as spleen (x-axis) versus MLN (y-axis). Red dashed indicates 1:1 ratio MLN:Spleen uptake. Linear regression for data shown as black line. (G). Graph showing pooled data of the ratio of spleen and MLN uptake for each immune population comparing cpPA and cpAA (H). Data is representative (A,B), mean +/- SEM (C,D,F), individual mice (G), and individual ratios plus median (H); for 3 (A-D,F), 6(E) and 4-5 (G-J) mice from at least 2 individual experiments. Statistical analysis was performed using ANOVA or non-parametric tests as appropriate; Šidák (C,D,E), Tukey (F,H) post-tests, linear regression (I) or a Mann Whitney test (J). ns, not significant; *, p < 0.05; **, p < 0.01; ***, p < 0.001; ****, p < 0.0001.

Importantly, beyond this global reduction, we observed striking differences in the relative uptake profiles across immune cell populations that were specific to cpOA. These differences were not predicted by intrinsic uptake capacity measured ex vivo and therefore suggested that access to cpOA is unequally distributed within the splenic microenvironment. Discrepancies emerged between B cells and T cells, as well as among myeloid populations, including cDC1, cDC2, and macrophages (Fig. 6A). *Ex vivo*, when spatial constraints were removed, B cells exhibited a greater intrinsic capacity for cpOA uptake than other lymphocytes, including T cells and NK cells (Fig. 6A). Similarly, cDC2 cells showed higher cpOA uptake capacity than cDC1s under *ex vivo* conditions (Fig. 6A). In contrast, these intrinsic hierarchies were not reflected *in vivo*. Instead, T cells acquired cpOA as efficiently as, or more efficiently than, B cells, and cDC1s exceeded cpOA uptake by cDC2s within the intact spleen (Fig. 6A).

To directly compare intrinsic uptake capacity with nutrient acquisition in vivo, the ratio of in vivo to ex vivo uptake was calculated for each immune-cell population (Fig. 6C,D). For cpOA, substantial differences were observed between immune subsets. B cells, cDC2s and macrophages retained a markedly smaller fraction of their ex vivo uptake capacity in vivo than T cells and cDC1s (Fig. 6C), indicating that intrinsic FA uptake capacity did not accurately predict FA acquisition within the intact spleen. In contrast, the in vivo:ex vivo ratios for HpG were comparatively uniform across immune populations (Fig. 6D), indicating that amino acid acquisition measured in vivo more closely reflected the intrinsic transport capacity revealed under ex vivo conditions. These findings suggest that access to circulating FA is more strongly influenced by local tissue constraints than access to circulating amino acids.

Having established that multiplexed in vivo nutrient-tracing could reveal differences in nutrient acquisition within a given tissue, we next asked differences in nutrient accessibility across tissues could also be detected. We focused on the spleen and thymus because these organs differ markedly in their vascular architecture and exposure to circulating nutrients. The spleen is a highly perfused organ with an open sinusoidal vasculature that directly samples circulating blood, whereas the thymus possesses a compartmentalized low-permeability vasculature that forms the blood-thymus barrier and restricts access to circulating factors ^28^. Importantly, T cells isolated from both organs have the capacity to take up cpOA and HpG ex vivo (Fig. 1)^13^, indicating that both populations possess the intrinsic machinery required to acquire these nutrients. To provide the most stringent test of nutrient accessibility, we focused on DN3 and DN4 thymocytes, developmental stages characterized by high metabolic activity and elevated cpOA uptake capacity (Fig. 1K). While in vivo cpOA uptake was detected in splenic T cells (Fig. 5), there was no cpOA detection in DN thymocytes (Fig. 6E). In contrast, HpG uptake was observed in both splenic T cells and thymocytes (Fig. 5, 6E). To interrogate whether these results were due to temporal differences in cpOA delivery to the thymus versus the spleen, a time course experiment was performed where spleens and thymuses were isolated at different timepoint post i.v. injection of cpOA and HpG. Splenic CD4 and CD8 T cells exhibited in vivo cpOA uptake, with a clear signal above PBS controls, from 5 to 120 min post injection (Fig. 6F). In contrast, no cpOA signal was detected in DN3 or DN4 thymocytes at any time point interrogated (Fig. 6F). Thus, despite retaining the intrinsic capacity to acquire FA, DN3 and DN4 thymocytes lacked detectable access to circulating cpOA in vivo, whereas access to circulating amino acids was preserved (Fig. 6E,F).

Taken together, these data demonstrate that cpOA uptake *in vivo* is not determined solely by cell-intrinsic uptake capacities but is strongly shaped firstly by tissue architecture and cellular positioning within the spleen (B cells versus T cells), and secondly by the type of tissue in question (Spleen vs Thymus). In contrast, HpG uptake closely mirrors intrinsic Slc1a5-mediated transport potentials with splenocytes and is more broadly distributed between tissues. The simplest explanation for the divergence within the spleen is that the microanatomical organization of the spleen differentially influences access of immune-cell subsets to circulating FA. Consistent with this idea, cpOA circulates as part of a large albumin-bound complex that is expected to experience greater spatial constraints on tissue distribution than the freely diffusible amino acid probe HpG. The size of FA-albumin complexes is likely the reason why cpOA does not access the thymus as potential antigenic proteins from the blood must be excluded from thymus where T cell selection is occurring.

### Fatty-acid-specific systemic distribution impacts immune-cell access *in vivo*

Having discovered how cpOA distribution and cell acquisition in vivo is not predicted by immune uptake capacities, we next interrogated whether different FA species distribute differently *in vivo.* Representative saturated (cpPA), monounsaturated (cpOA), and polyunsaturated (cpAA) FA were i.v. injected separately into mice. Uptake was assessed in the spleen and mesenteric lymph nodes (MLN), two immune tissues that differ fundamentally in their exposure to intravenously delivered metabolites (Fig. 6G). The spleen is directly perfused by systemic circulation and receives high blood flow, whereas the MLN primarily samples nutrients and metabolites delivered via lymphatic drainage. Strikingly, despite cpAA consistently exhibiting the highest uptake capacity *ex vivo* across all models examined (Fig. 1, 2, 4), this hierarchy was reversed *in vivo*. Following intravenous delivery, cpAA uptake by immune cells in both the spleen and MLN was significantly lower than uptake of the saturated fatty acid cpPA across both lymphoid and myeloid subsets (Fig. 6G). Moreover, cpPA showed significantly greater incorporation into immune cells than the monounsaturated cpOA (Fig. 6G), indicating pronounced FA-species specific differences in systemic transport and tissue access *in vivo*.

To directly assess fatty-acid transport between tissues, mean fluorescence intensity (MFI) values for each immune subset were plotted as paired spleen versus MLN uptake, generating an x-y relationship in which samples lying along the x = y line indicates equivalent uptake in both tissues. cpPA uptake clustered closely along this line, demonstrating efficient transport and distribution of cpPA across murine tissues (Fig. 6H). In contrast, cpAA uptake values were consistently shifted below the x = y line, indicating substantially greater uptake in the spleen than in the MLN and suggesting impaired delivery of cpAA to more distal lymphoid sites. Consistent with this interpretation, the calculation of the spleen-to-MLN uptake ratio revealed values close to 1 for cpPA across immune subsets, whereas ratios for cpAA were significantly lower (Fig. 6I). Together, these analyses demonstrate with single cell resolution that cpPA is efficiently transported throughout the organism, while cpAA shows more limited distribution beyond highly perfused tissues. These findings highlight a critical distinction between *ex vivo* uptake assays and nutrient acquisition *in vivo*. While *ex vivo* assays reveal the intrinsic capacity of immune cells to take up FA, *in vivo* access is dominated by FA-specific systemic distribution, tissue perfusion, and microanatomical constraints.

### Cell identity and tissue environment exert distinct influences on nutrient uptake in vivo

The data above argue for an important role for tissue location in determining how much circulating FA an immune cell will acquire. We next wanted to directly test the relative contributions of cell identity and tissue environment to the nutrient acquisition in lymphocytes in vivo. Therefore, we simultaneously measured uptake of the clickable FA cpOA and the amino acid probe HpG into lymphocyte in diverse tissues following systemic administration (Fig. 7A). Lymphocytes isolated from blood, spleen, liver, lung, and mesenteric and axillary lymph nodes (MLN and ALN) were analysed by flow cytometry for cpOA uptake, HpG uptake and expression of CD98, a broadly expressed nutrient transporter-associated protein widely used as a surrogate marker of lymphocyte metabolic activation and nutrient uptake capacity. To identify patterns of nutrient acquisition independently of predefined immune-cell classifications dimensionality reduction was performed using cpOA uptake, HpG uptake, CD98 expression and the morphological parameters FSC(A) and SSC(A), while lineage markers were excluded from UMAP generation and subsequently overlaid onto the resulting map.

**Figure 7.**
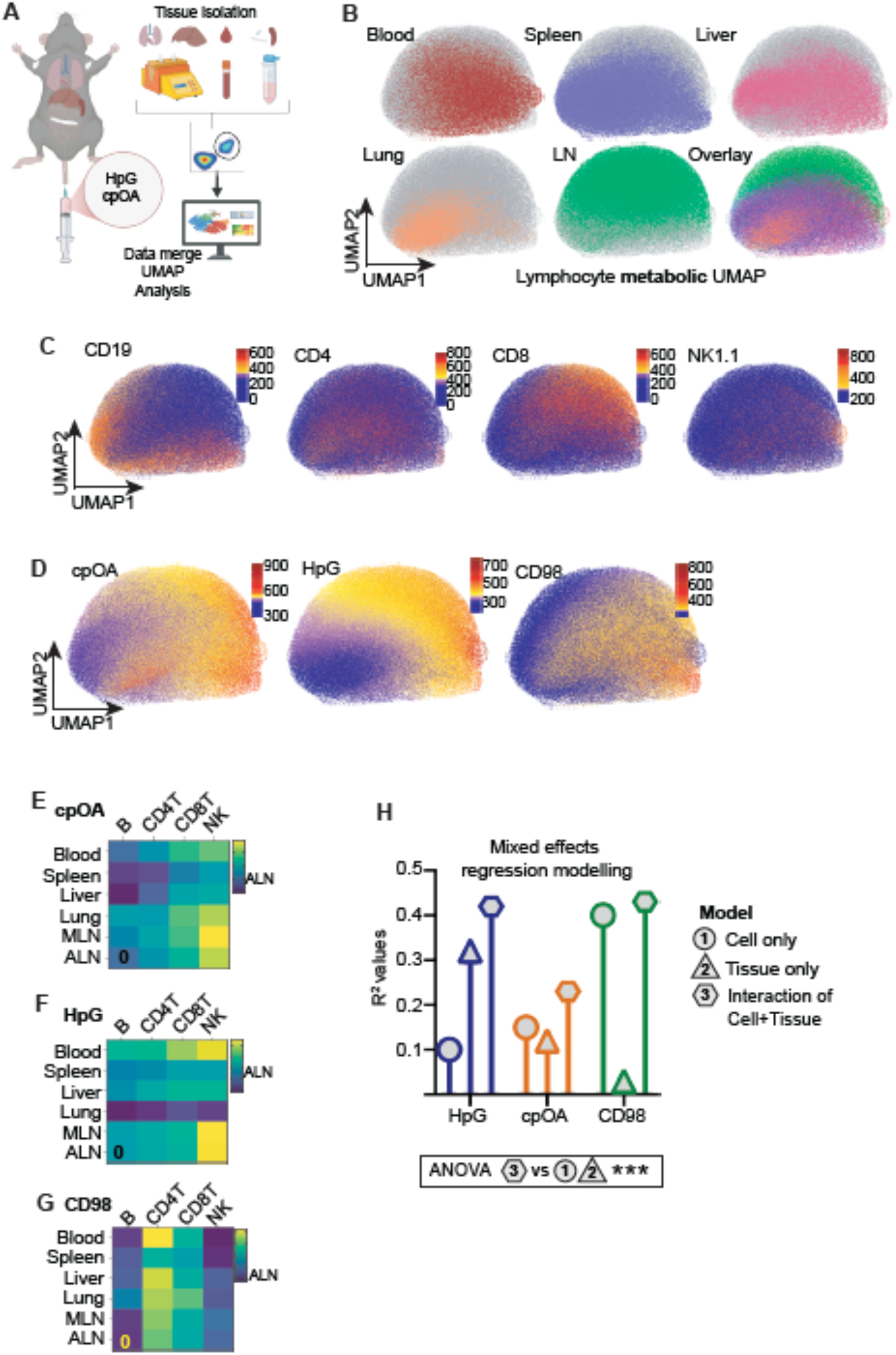
Tissue environment and cellular identity differentially shape nutrient acquisition in vivo. **(A-D)** Mice were i.v. injected with 10 mg/kg cpOA, 100 mg/kg HPG and tissues harvested after 30 min (Spleen, Lung, Liver, Blood, ALN, MLN) and prepared for flow cytometry analysis by staining with cell lineage markers and CD98. Flow data was filtered to include only live lymphocytes (T cells (CD4^+^ and CD8^+^), B cells and NK cells) and data exported, combined from 3 separate mice and cellular relationships visualised using a UMAP (A). The distribution of the different tissues in the UMAP is visualised (B) and the cell identity markers are overlaid onto the UMAP (C). The intensity of in vivo cpOA uptake, in vivo HpG uptake, CD98 expression is shown per cell in the UMAP (D). Mean data estimated by mixed-effect modelling for cpOA (E) and HpG uptake (F) and CD98 expression (G) is shown in heatmaps for each lymphocyte and each tissue, normalised to ALN. (H) R^2^ values for each mixed-effects regression indicate the proportion of variation explained by each model. An ANOVA test was used to compare the fit of the models to the data and show that that model 3, the interaction of cell type and tissue location, fitted the data significantly better than model 1 and 3. ***, p < 0.001.

Visualisation of cells according to tissue origin revealed distinct tissue-enriched regions within the metabolic UMAP space (Fig. 7B). Overlay of lineage markers demonstrated that B cells occupied a discrete region of the UMAP despite not contributing to its generation, indicating that B cells possess a distinct combination of nutrient-uptake, transporter-expression and morphological features compared with other lymphocyte populations (Fig. 7C). In contrast, CD4 T cells, CD8 T cells and NK cells showed substantial overlap, suggesting greater similarity in these parameters. Mapping of cpOA uptake, HpG uptake and CD98 expression onto the same UMAP revealed distinct metabolic gradients across the landscape, indicating considerable metabolic heterogeneity among lymphocytes in vivo (Fig. 7D).

To further dissect the relative influence of tissue location and lymphocyte identity, data were summarized by tissue and cell type. Inspection of the heatmaps revealed distinct organizational features for each metabolic parameter. HpG uptake displayed a strong tissue-associated pattern that was largely conserved across lymphocyte populations, with all subsets exhibiting elevated uptake in blood and reduced uptake in lung (Fig. 7E).

In contrast, CD98 expression followed a predominantly cell-type-associated pattern, with CD4 T cells consistently expressing the highest levels of CD98 across tissues (Fig. 7F). cpOA uptake exhibited a distinct intermediate profile. NK cells consistently displayed the highest cpOA uptake yet differences between tissues were comparatively modest and varied across lymphocyte populations (Fig. 7G). Thus, visual inspection suggested that HpG uptake was organized primarily by tissue location, CD98 expression by lymphocyte identity, and cpOA uptake by contributions from both factors.

These observations were quantified using mixed-effects modelling (Fig. 7H). In this analysis, the R^2^ value represents the proportion of variation that can be explained by the variables included in the analysis: cell type (model 1), tissue location (model 2) and the interaction of cell types and tissue locations (model 3). Tissue location explained substantially more variation in HpG uptake than lymphocyte identity (R^2^ = 0.32 versus 0.10), while inclusion of both variables increased the explanatory power of the model (R^2^ = 0.42). This indicates that amino acid acquisition is strongly influenced by the local tissue environment. In contrast, cell identity and tissue location explained similar proportions of the variation in cpOA uptake (R^2^ = 0.15 and 0.12, respectively) and together accounted for only a modest fraction of the overall variability (R^2^ = 0.23). Thus, cpOA uptake was influenced by both lymphocyte identity and tissue location, but neither factor alone strongly predicted cpOA acquisition, suggesting additional sources of heterogeneity contribute to cpOA uptake in vivo. CD98 expression displayed a strikingly different behaviour, with lymphocyte identity explaining most of the variation (R^2^ = 0.41) and tissue location contributing very little (R^2^ = 0.03). Addition of tissue information produced almost no improvement in model performance (combined R^2^ = 0.43), indicating that CD98 expression is determined predominantly by intrinsic differences between lymphocyte populations rather than their tissue environment.

Collectively, these findings demonstrate that the relative contributions of tissue environment and cell identity differ substantially between metabolic pathways in vivo. Amino acid uptake is strongly associated with tissue location, CD98 expression is primarily linked to lymphocyte identity, and FA uptake reflects contributions from both factors while remaining comparatively heterogeneous within tissues.

## Discussion

FA are essential nutrients that support immune-cell bioenergetics, membrane synthesis, and signalling; however, existing approaches for measuring FA uptake at single-cell resolution have important limitations. Here, we establish cyclopropene-tagged FA (cpFAs) as minimally perturbing probes for quantifying FA uptake in immune cells. Consistent with physiological protein-mediated uptake, cpFA incorporation was temperature dependent, competitively inhibited by native FA, and influenced by established FA handling pathways. In contrast, the widely used probe BODIPY-C16 accumulated independently of temperature and failed to report active FA transport, further supporting concerns that bulky fluorophore-conjugated metabolites do not necessarily reflect physiological nutrient uptake^29^. These findings reinforce a general principle emerging across nutrient-uptake studies: direct conjugation of large fluorophores can substantially alter nutrient transport, whereas bioorthogonal approaches preserve native uptake mechanisms by separating transport from fluorescent detection^13,14^.

FA uptake is increasingly understood as a coordinated process involving membrane translocation, intracellular trapping, and metabolic utilization rather than a simple transport event ^30–32^. Consistent with this model, we identify FABP5 as a major determinant of cpFA uptake in immune cells and demonstrate a more selective role for CD36 in myeloid populations. The strong ability of polyunsaturated FA to compete with cpOA uptake further supports the conclusion that cpFAs engage endogenous FA handling pathways rather than undergoing nonspecific membrane partitioning ^19,20,33–36^. Together, these data indicate that cpFA uptake reflects the integrated activity of physiological transport and intracellular lipid-handling machinery.

Application of the cpFA platform revealed substantial heterogeneity in FA uptake across immune populations and tissues. Regulatory T cells displayed enhanced FA uptake relative to naïve CD4 T cells, with the most pronounced phenotype observed in the small-intestinal lamina propria. Importantly, naïve CD4 T cells exhibited relatively similar uptake capacities across tissues, whereas effector-memory and Treg populations showed marked tissue-specific differences. These findings suggest that metabolic adaptation to local environments is acquired during differentiation and tissue residency rather than representing a general property of all CD4 T cells^37,38^. The elevated uptake of multiple FA classes by intestinal Tregs is consistent with adaptation to a nutrient-rich environment continuously exposed to dietary lipids and microbiota-derived metabolites and highlights FA acquisition as a previously underappreciated aspect of tissue-specific Treg biology^37,39^.

A central insight arising from this work is the distinction between intrinsic uptake capacity and nutrient accessibility in vivo. Although arachidonic acid displayed the highest uptake capacity across multiple immune populations ex vivo, it exhibited relatively limited distribution following intravenous administration. In contrast, palmitate showed lower intrinsic uptake but substantially broader tissue accessibility. These observations indicate that nutrient acquisition in vivo is determined not only by cellular transport and metabolic capacity but also by nutrient-specific differences in systemic distribution, clearance, and tissue^40^ delivery ^40,41^. AA is a bioactive FA and is removed rapidly from the blood into tissues and incorporated into cellular membranes, whereas PA is handled as a well-buffered, stable circulating pool for inter-organ transport. Consequently, measurements made under uniform ex vivo conditions cannot fully predict nutrient uptake within intact organisms. Consequently, measurements made under uniform ex vivo conditions do not predict nutrient availability within intact organisms; rather these are a readout for the capacity of the cell to take up the nutrient, should it be available. These data suggest that tissue location and vascular access are key determinants of immune-cell FA acquisition *in vivo*, with important implications for tissue-specific immune niches as well as pathological contexts such as tumors and infection.

Our in vivo analyses further reveal how tissue architecture shapes access to circulating nutrients. Within the spleen, uptake patterns differed markedly from intrinsic ex vivo uptake capacities, suggesting that blood flow, cellular positioning, and competition among neighbouring cell populations influence which cells gain access to circulating FA.

An even more striking example was observed in the thymus, where a blood-thymus barrier exists and uptake of circulating cpOA was essentially absent despite robust ex vivo uptake capacity ^28^. In contrast, metabolically active DN3 and DN4 thymocytes readily acquired the systemically administered amino acid analogue HpG. These findings highlight that nutrient access across tissue barriers is selective rather than absolute and establish the blood-thymus barrier as a previously unrecognized regulator of systemic lipid availability. Indeed, local production of OA was shown to be essential for thymic Treg differentiation ^17^. More broadly, they illustrate how tissue-level organization can impose metabolic constraints independently of cell-intrinsic uptake machinery^5,6^.

The ability to simultaneously measure FA and amino acid uptake provided an opportunity to directly compare the relative contributions of tissue environment and cellular identity to nutrient acquisition. Multiparameter analysis revealed that these contributions differ markedly between metabolic pathways. HpG uptake exhibited a strong tissue-dependent signature across lymphocyte populations, with tissue location explaining substantially more variation than lymphocyte identity. In contrast, expression of the nutrient transporter-associated protein CD98 was explained predominantly by lymphocyte identity and showed relatively little dependence on tissue location. FA uptake occupied an intermediate position, with both tissue environment and cell identity contributing to cpOA acquisition. However, these factors together explained only a modest proportion of the overall variation in cpOA uptake. This observation suggests that FA acquisition is influenced by additional sources of heterogeneity not captured by broad tissue or lineage classifications.

These findings extend the central conclusion of this study that nutrient accessibility represents a distinct layer of metabolic regulation in vivo. While amino acid uptake could be predicted to a considerable extent from tissue location, FA uptake remained comparatively heterogeneous even among cells of the same lineage within the same tissue. Such variability is consistent with our observations that FA accessibility is shaped by anatomical barriers, vascular organization and local nutrient distribution. Consequently, tissue identity alone may be insufficient to predict FA availability, implying that metabolic niches operate at a finer spatial scale than whole-organ classification.

Importantly, direct nutrient uptake measurements revealed patterns that would not have been predicted from expression of a commonly used metabolic markers, such as CD98. Although CD98 expression was largely associated with lymphocyte identity, amino acid and FA uptake displayed substantially different relationships with tissue environment. These findings argue that immune-cell metabolism cannot be represented by a single metric but instead reflects multiple layers of regulation that are differentially influenced by intrinsic cellular programming and local environmental conditions. Collectively, these results highlight the value of direct in vivo nutrient-tracking approaches for resolving how tissue context shapes immune-cell metabolism.

Beyond FA biology, the bioorthogonal strategy described here provides a general framework for investigating nutrient accessibility in vivo. Because substrate uptake is uncoupled from fluorophore detection, the same principle should be applicable to diverse classes of metabolites whose transport may be distorted by fluorescent conjugation ^12,13,29^. Furthermore, the successful simultaneous measurement of cpOA and HPG uptake highlights the future potential for multiplexed in vivo nutrient uptake measurements using orthogonal click-chemistry reactions. Such approaches should enable direct mapping of nutrient accessibility across tissues and provide new opportunities to determine how vascular organization, tissue architecture, and local metabolic niches influence cellular metabolism in vivo.

An important consideration when interpreting in vivo nutrient-uptake measurements is that reduced incorporation of a clickable nutrient does not, by itself, define the mechanism responsible for the observed phenotype. Limited labeling may reflect restricted physical access of the nutrient to a particular tissue niche, competition for nutrient acquisition by neighbouring cells, or competition from high concentrations of endogenous unlabeled nutrient within the local microenvironment. These mechanisms are not mutually exclusive and may operate simultaneously within complex tissues. Consequently, the approach described here should be viewed primarily as a means of identifying cellular and anatomical niches that differ in nutrient accessibility, rather than an absolute measure of nutrient availability or transporter activity alone. Combining bioorthogonal nutrient-uptake measurements with spatial imaging, metabolomics or perturbation of candidate transport pathways should provide a powerful strategy for resolving the mechanisms underlying these differences.

Together, these findings establish bioorthogonal nutrient-uptake profiling as a powerful approach for identifying cellular and anatomical niches in which nutrient accessibility shapes immune-cell metabolism in vivo.

## Author contributions

Conceptualization: X.W., G.H., L.V.S., M.v.d.S., S.I.v.K., D.K.F. Methodology: X.W., G.H., L.R., L.I.B, S.I.v.K., D.K.F. Investigation: X.W., G.H., C.C., C.L., K.S., K.B., M.B., J.M.P., W.P.F.D. Writing – Original Draft: X.W., G.H., C.C., S.I.v.K, D.K.F. Writing – Review & Editing: All authors. Funding Acquisition: B.E., M.v.d.S., S.I.v.K., D.K.F. Supervision: B.E., S.I.v.K., D.K.F.

## Acknowledgements

D.K.F lab is supported by Taighde Éireann/Research Ireland [22/FFP-A/10326 and IRCLA/2023/1402]. SIvK was funded by the ERC Consolidator Grant (grant number 865175) and by the Institute for Chemical Neuroscience (iCNS: Gravitation grant 024.006.009) of the Dutch Research Council (NWO). MvdS was funded by the Netherlands Organization for Scientific Research VICI-grant (grant number 724.017.002) and by the Institute for Chemical Neuroscience. B.E. lab is supported by Dutch Research Council (NWO) [VIDI Grant 91719349 and OCENW.M.23.183]. We thank the Flow Cytometry Core Facility, Microscopy and Imaging Centre and Comparative Medicine Unit at Trinity College Dublin for technical support.

## Data and Code Availability

All data supporting the findings of this study are available from the corresponding authors upon reasonable request. Custom analysis scripts are available at GitHub: github.com/changliu979/nutrient-uptake-in-vivo

## STAR METHODS

### Mice

In Trinity College Dublin, Female C57BL/6J WT mice were obtained from Envigo or bred in house and C57BL/6J CD36-/- mice were bred in house. All mice were used for experiments between 6 and 12 weeks of age. Mice were housed under 12:12 light cycle in a relative humidity of 45–65% and a temperature between 20℃ and 24℃. Mouse experiments were approved by and in compliance with the Irish Health Products Regulatory Authority (Project license AE19136/P177) and the Animal Research Ethics Committee (AREC) at Trinity College Dublin.

At the Leiden Institute of Chemistry (LIC), C57BL/6J male and female mice were bred under specific pathogen free (SPF) conditions. Animal license number AVD1160020198832. Mice between 6 weeks and 6 months old were culled by cervical dislocation. Animal experiments were approved by the Dutch Central Authority for Scientific Procedures on Animals (CCD) and performed in accordance with European Union Directive 2010/63EU, Recommendation 2007/526/EC and local government regulations.

### Cell culture

NK92MI human natural killer cells were maintained in RPMI-1640 medium supplemented with 12.5% (v/v) fetal bovine serum (FBS), 12.5% (v/v) horse serum, 2 mM glutamine, 0.2 mM inositol, 0.02 mM folic acid, 0.1 mM β-mercaptoethanol, and 100 U/mL penicillin–streptomycin. HEK293T human embryonic kidney cells were cultured in DMEM high glucose medium containing 10% (v/v) FBS and 100 U/mL penicillin– streptomycin. THP-1 human monocytic leukemia cells were maintained in RPMI-1640 medium supplemented with 10% (v/v) FBS and 100 U/mL penicillin–streptomycin. All cell lines were incubated at 37°C in a humidified atmosphere with 5% CO₂.

### FABP5 knock out in NKG2MI and THP-1 cell lines

FABP5 knockout cell lines were generated using CRISPR/Cas9-mediated genome editing. The sgRNA targeting exon 1 of the human FABP5 gene (5′-AAGGAGCTAGGTGAGGCACC-3′) was designed using the CRISPOR online tool (http://crispor.tefor.net/). NK92MI and THP-1 cells were cultured in fresh complete medium overnight prior to electroporation. For each sample, 2.5×106 cells were collected and washed three times with cold PBS (without Ca2+and Mg2+). Cell pellets were incubated on ice for 5 min and resuspension in 100 μL electroporation buffer. Recombinant S. pyogenes Cas9 protein (100 pmol) and FABP5 sgRNA (100 pmol) were preassembled into ribonucleoparticle (RNP) complex and mix with 3 μg linear dsDNA neomycin knockout template in 100 μL electroporation buffer. The cell suspension and RNP/donor DNA mixture were gently mixed and transferred into sterile 0.2 cm electroporation cuvettes. Electroporation was performed using cell type specific programs recommended for the Lonza® Nucleofector® II electroporation system according to previously published protocols (STAR Protocols, 2024, 5:103123). After electroporation, 1 mL of pre-warmed complete culture medium was gently added to the cuvette without disturbing the cells. Cells were allowed to recover in a humidified incubator for 15 min before transferring into T75 flasks containing 5 mL complete medium. Cells were maintained for 2–3 days until reaching approximately 80–90% confluency. To enrich successfully transfected cells, G418 selection was performed at concentrations of 250 μg/mL for THP-1 cells and 1 mg/mL for NK92MI cells for 7 days. Surviving cells were subsequently maintained in culture medium supplemented with a lower concentration of G418 (10 μg/mL). For single cell cloning, cells were diluted to a density of 0.5 cells per 100 μL and seeded into 96-well plates with 200 μL per well. Individual single cell colonies were expanded after reaching approximately 80% confluency and subsequently transferred into 24-well plates for further validation and downstream experiments.

### Isolation of Immune cells from tissues

Spleen, thymus, liver, axillary lymph nodes (ALN), and mesenteric lymph nodes (MLN) were harvested from C57BL/6J WT mice or CD36-/- mice and mechanically dissociated using the plunger end of a syringe. Tissue suspensions were digested in 5 mL RPMI-1640 medium with 1 mg/mL collagenase D and 2000 U/ml DNase I for 20 minutes at 37°C in a 5% CO₂ incubator. Lungs were finely minced and digested in 3 mL RPMI-1640 medium containing 1 mg/mL collagenase D and 2000 U/mL DNase I. Tissue samples were incubated for 30 min at 37°C with agitation at 180 rpm. Small intestines were carefully cleared of surrounding fat, opened longitudinally, and gently cleaned to remove feces and mucus. Intestinal tissues were washed in 20 mL Ca²⁺/Mg²⁺-free HBSS containing 2 mM EDTA and cut into 1–2 cm pieces. To remove epithelial cells, tissues underwent three rounds of washing in HBSS containing 2 mM EDTA, with incubation for 15 min at 37°C with agitation at 180 rpm during each round. Following epithelial removal, tissues were digested in 10 mL RPMI-1640 medium containing 1 mg/mL collagenase D and 2000 U/mL DNase I for 20 min at 37°C with agitation at 180 rpm. Following enzymatic digestion, the resulting cell suspension was passed through 70–100 µm sterile strainers and centrifuged. Red blood cells were lysed using red blood cell lysis buffer for 5 minutes at room temperature. Cells were subsequently washed with fresh medium, centrifuged at 1500 rpm for 5 minutes, and resuspended in fresh medium for downstream experiment. Peripheral blood was collected into EDTA-coated tubes and diluted in 1mL PBS. Red blood cells were lysed using red blood cell lysis buffer for 10 min at room temperature. Samples were centrifuged at 400 g for 5 min and washed twice with PBS supplemented with 2% FBS. Cells were subsequently resuspended in staining buffer for downstream experiment.

### T cells enrichment from tissues

Immune cell suspensions isolated from different tissues (Spleen, Lung, MLN, and SILP) were stained separately with anti-CD45 antibodies conjugated to distinct fluorophores for 15 min at room temperature as indicated in Figure4A. Cells were then washed and pooled on a per-mice basis to balance the relative abundance of T cells across tissues. For each mouse, the pooled sample consisted of 5 × 10⁶ lung cells, 5–10 × 10⁶ SILP cells, 1 × 10⁶ spleen cells, and 1 × 10⁶ MLN cells. Total CD3⁺ T cells were subsequently isolated from the pooled cell suspension by magnetic negative selection according to the manufacturer’s instructions.

### In vivo NK cells activation

Poly(I:C) was used to activate the splenic NK cells in vivo as previously described (Ref). Briefly, Poly(I:C) was diluted in PBS and injected i.p. at a total dose of 200 µg per mice for 18h. Equal volume of PBS were injected to control mice as vehicle control. On the second day, the mice were culled, and the spleens processed as described above.

### Cyclopropane-tagged fatty acid (cpFA) uptake

Cell lines and immune cells from tissue were collected and pre-stained with LIVE/DEAD Fixable Dead Cell Stain for 30 minutes at room temperature in the dark. An appropriate number of cells were then seeded into V-bottom 96-well plates, washed twice with HBSS, and resuspended in 100 µL of HBSS.

In parallel, HBSS medium containing twice the indicated concentration of cpFA (2× cpFA solution) was prepared and either pre-warmed in a 37°C incubator or chilled on ice (4°C, cold controls) for 20 min to allow temperature equilibration. Next, 100 µL of the 2× cpFA solution was added to the cells, and the plates were returned to the incubator or ice for the designated incubation period. After incubation, cells were centrifuged, washed once with PBS, and fixed with 200 µL of 1% paraformaldehyde for 30 min at room temperature in the dark. Plates were centrifuged again, supernatants were removed, and cell pellets were resuspended in PBS and kept at 4°C for subsequent click-chemistry labeling and flow-cytometry staining.

To assess whether different fatty acids competitively inhibited cpOA uptake, the indicated concentration fatty acids were added to uptake medium containing 2× concentrated cpOA, and uptake assays were performed as described above.

### Bioorthogonal amino acid (HPG) uptake assay

The bioorthogonal amino acid uptake assay using L-homopropargylglycine (HPG) was performed as previously described ^13^. Briefly, cells from tissue were collected and pre-stained with LIVE/DEAD Fixable Dead Cell Stain for 30 minutes at room temperature in the dark. Cells were then washed twice with HBSS and resuspended in 100 µL of pre-warmed HBSS in a 37°C incubator for 20 min to allow temperature equilibration. Next, 100 µL of 800 µM HPG-HBSS solution was added to the cells, and cells were returned to the incubator for the designated incubation period. After incubation, cells were centrifuged, washed once with PBS, and fixed with 200 µL of 1% paraformaldehyde for 30 min at room temperature in the dark. Plates were centrifuged again, supernatants were removed, and cell pellets were resuspended in PBS and kept at 4°C for subsequent click-chemistry labeling and flow-cytometry staining.

### Click chemistry

Cells were permeabilized for 20 min at room temperature using 0.01% saponin in PBS. After permeabilization, the plate was centrifuged, cells were washed twice with PBS. For the cpFA uptake assay, cell pellets were resuspended in 30 µL PBS with 1µM Tetrazine-BODIPY FL for 45 min at room temperature in the dark. For the HPG uptake assay, cell pellets were resuspended in 30 µL click-mixture solution (1 mM CuSO4, 10 mM NaAsc, 1mM THPTA, 10 mM Aminoguanidine in PBS) with 1uM AZDye 647 azide for 45min at room temperature in the dark. After the click reaction, cells were centrifuged, supernatants were removed, and cell pellets were washed twice with PBS and once with FACs buffer, and cell pellets were resuspended in PBS and kept at 4°C for antibody staining or flow cytometer.

### BODIPY-C16 uptake assay

NK92MI cells were pre-stained with LIVE/DEAD Fixable Dead Cell Stain for 30 minutes at room temperature in the dark. An appropriate number of cells were then seeded into V-bottom 96-well plates, washed twice with uptake medium, and resuspended in 75 µL of uptake medium.

In parallel, uptake medium containing 4× BODIPY-C16 was prepared and either pre-warmed in a 37°C incubator or chilled on ice (4°C, cold controls) for 20 min to allow temperature equilibration. Next, 25 µL of 4× BODIPY-C16 was added to the cells, and the plates were returned to the incubator or ice for 10 min. After incubation, cells were centrifuged, washed once with PBS, and fixed with 200 µL of 1% paraformaldehyde for 30 min at room temperature in the dark. Plates were centrifuged again, supernatants were removed, and cell pellets were resuspended in PBS and analyzed by flow cytometry.

### Confocal microscopy

The cpOA uptake assay and BODIPY-C16 uptake assay in NK92MI cells were performed as described above. For the cpOA uptake assay, following the click reaction, cells were washed four times with PBS and resuspended in 50 µL PBS containing 1% fatty acid free-BSA and 1 µg/mL DAPI for 45 min at room temperature in the dark to stain nuclei. For the BODIPY-C16 uptake assay, following the fixation, cells were washed twice with PBS and resuspended in 50 µL PBS containing 10 μg/mL Hoechst 33342 nuclear stain for 30 min at room temperature in the dark to stain nuclei. After nuclear stain, cells were centrifuged, washed twice with PBS, and resuspended in CyGEL™. The cell suspension was pipetted directly onto high-resolution microscope slides and allowed to solidify for 10 minutes at room temperature in the dark. Images were acquired using a Leica SP8 scanning confocal microscope. Quantification of mean pixel intensity was performed using IMARIS software by generating a pixel mask for the green fluorescence channel after applying an appropriate threshold to eliminate background fluorescence.

### Flow cytometry

Cells were pre-incubated with monoclonal antibody 2.4G2 (anti-mouse CD16/CD32 mAb) to block Fcγ receptors. Cells were then stained with fluorochrome-conjugated monoclonal antibodies for 45 minutes at room temperature in the dark. Acquisition was done on a BD FACSCanto II or BD LSRFortessa. Analysis, including dimensionality reduction by t-distributed stochastic neighbor embedding (tSNE), was done using FlowJo.

### Western Blot

WT and FABP5-/- NK92MI single clone cells were harvested at 1 × 106 cells per sample and washed twice with cold PBS prior to lysis in RIPA buffer supplemented with protease inhibitor cocktail on ice for 30 min. Equal amounts of protein were denatured by boiling, resolved by SDS–PAGE using precast gels, and transferred onto PVDF membranes. Membranes were blocked in 5% skim milk prepared in TBST (Tris-buffered saline containing 0.1% Tween-20) for 1 h at room temperature and incubated overnight at 4°C with primary antibodies against FABP5 and β-actin. On the second day, after three washes with TBST (15 min each), membranes were incubated with HRP-conjugated secondary antibodies for 1 h at room temperature. Following three additional washes with TBST, protein bands were visualized using enhanced chemiluminescence detection kit and imaged with Image Pro Plus. Band intensities were quantified using ImageJ software.

### In vivo administration of Clickable Nutrients

cpOA, cpAA, and cpPA were dissolved in DMSO to prepare an 85 mM stock solution, and HPG was dissolved in PBS to generate a 40 µg/µL stock solution. Both clickable nutrients were sterilized by filtration through a 0.22-µm filter. Mice were injected intravenously (i.v.) via the tail vein with the indicated doses of cpOA, cpAA, cpPA or/and HPG. Control mice received an equal volume of PBS as vehicle control. After the designated time points, mice were euthanized using CO₂ anesthesia followed by cervical dislocation. Different tissues were collected, and single-cell suspensions were prepared as described above. Cells were then subjected to surface phenotyping, fixation, and click chemistry according to the procedures described previously.

### Cell identity / tissue environment analysis for cpOA and HPG in vivo uptake

Data on the simultaneous uptake of cpOA and HPG by different immune cell populations across tissues were collected as described above. Single-cell measurements were analysed using linear mixed-effects model (lme4 version 2.0-1 package, effects version 4.2-5 package, R version 4.5.3), with cells treated as observations nested within individual mice. Separate models were fitted for each metabolic parameter, including HPG uptake, cpOA uptake, and CD98 expression. For each parameter, three models were evaluated. The first model included lymphocyte subset (B cells, CD4⁺ T cells, CD8⁺ T cells, and NK cells) as a fixed effect. The second model included tissue location (blood, spleen, liver, lung, MLN, and ALN) as a fixed effect. The third model included the fixed effects of lymphocyte subset and tissue location, together with their interaction, to determine whether the effect of tissue location differed among lymphocyte subsets. Mice identity was included as a random effect in all models to account for repeated measurements and baseline differences between individual animals. Model performance was assessed using the values of marginal R² and conditional R². Marginal R² represents the proportion of variance explained by the fixed effects alone, whereas conditional R² represents the proportion of variance explained by both the fixed and random effects.

### Statistical analysis

For experimental datasets analysed in GraphPad Prism, comparisons across multiple groups were performed using one way or two-way analysis of variance (ANOVA) with Tukey’s or Šídák post hoc tests, or mixed effects models (REML) where appropriate. Normality and homogeneity of variance were assessed using Shapiro–Wilk and Levene/Brown–Forsythe tests. When assumptions of parametric testing were not met, non-parametric alternatives were used as indicated. Outliers were excluded using a predefined criterion (±2.5 standard deviations from the mean). Unless otherwise stated, n indicates the number of mice; technical replicates were averaged per biological replicate prior to analysis. A p value <0.05 was considered statistically significant.

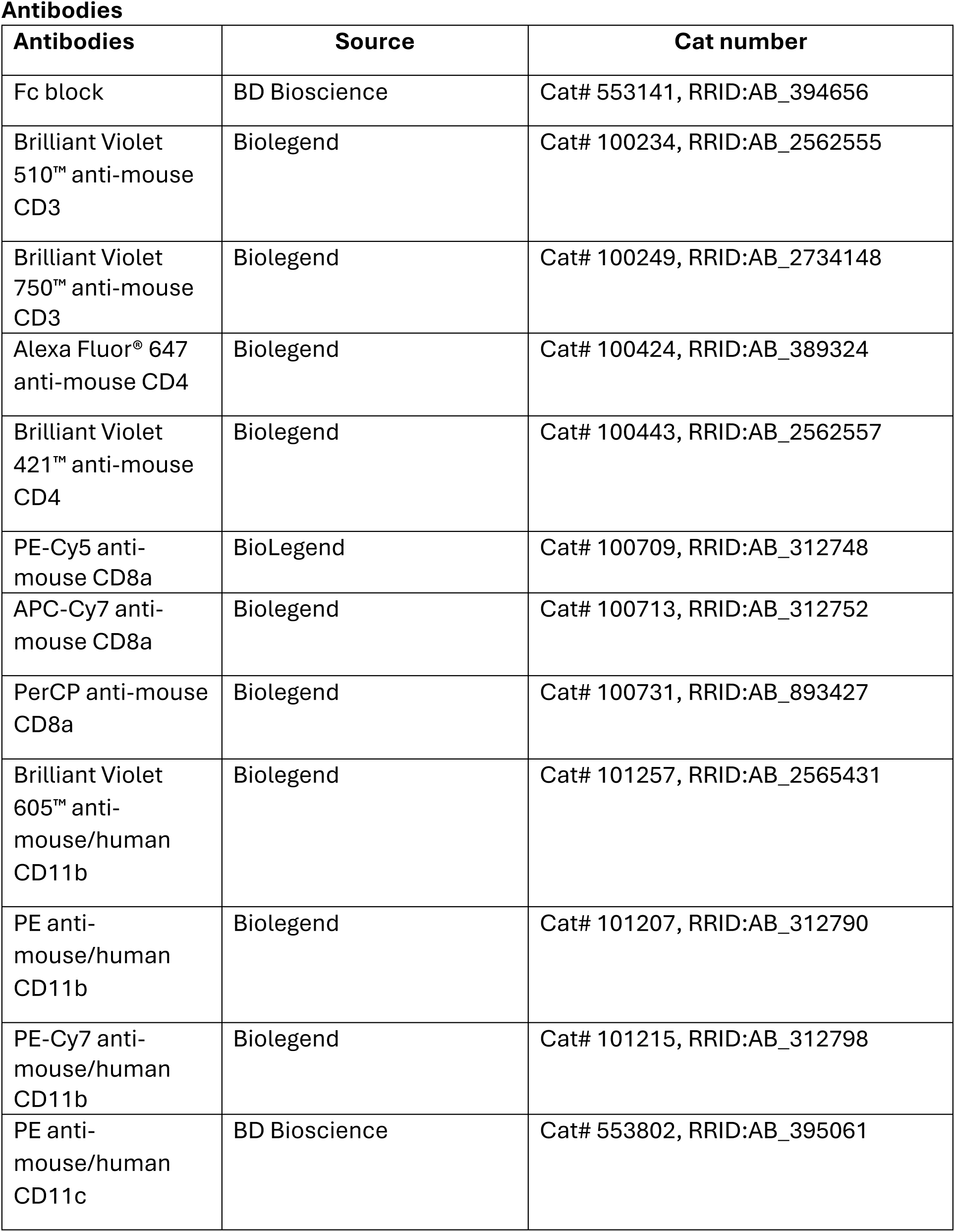
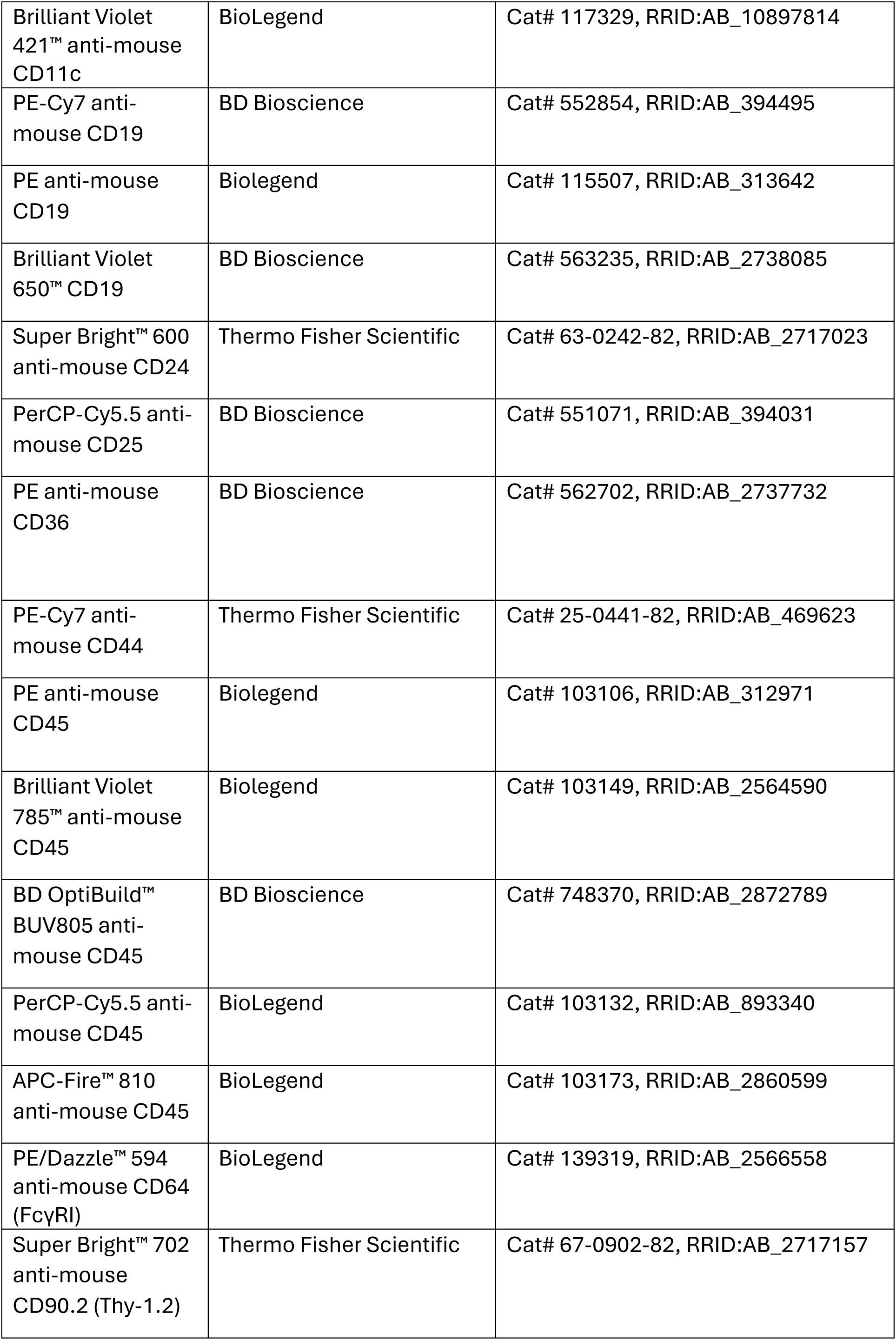
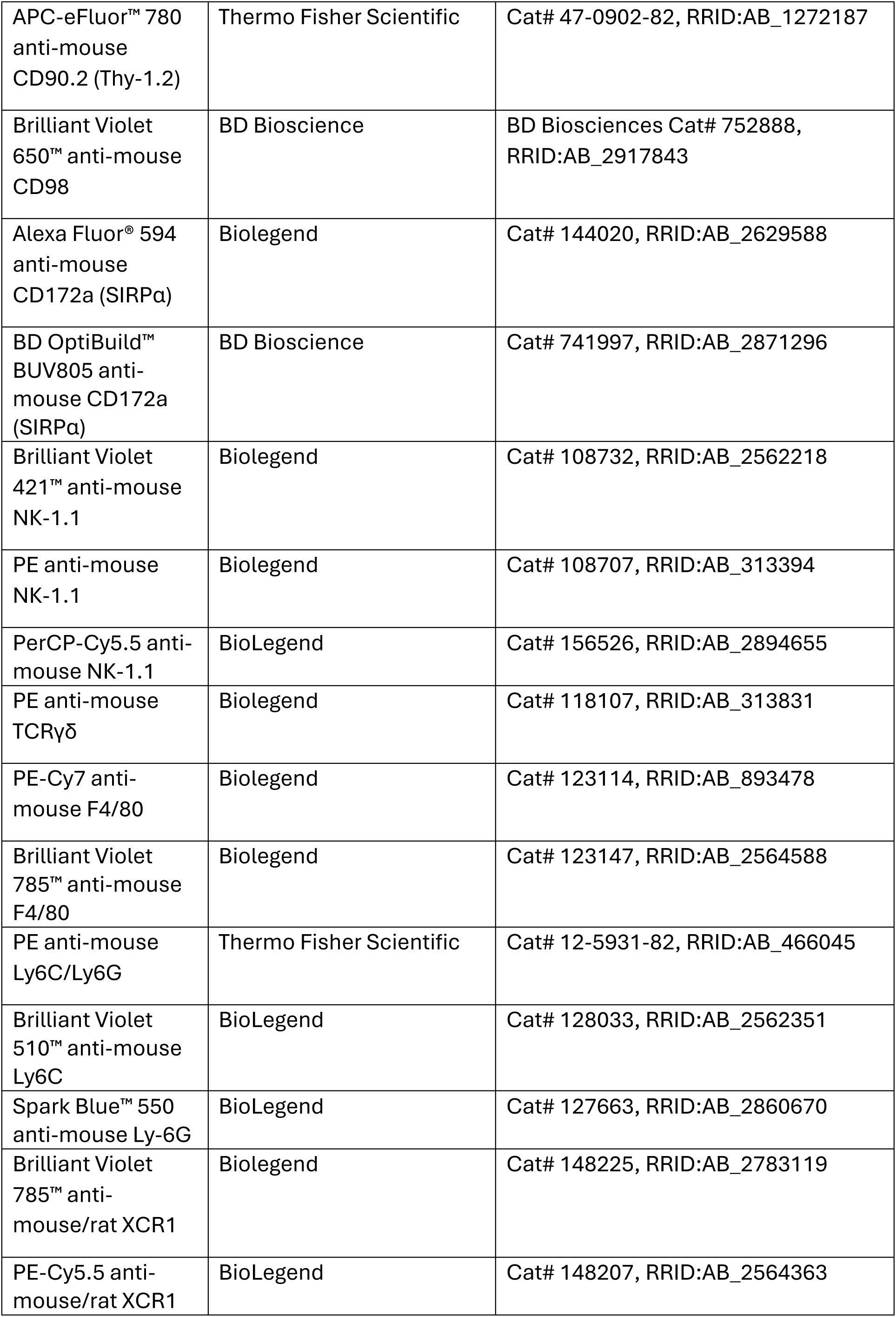
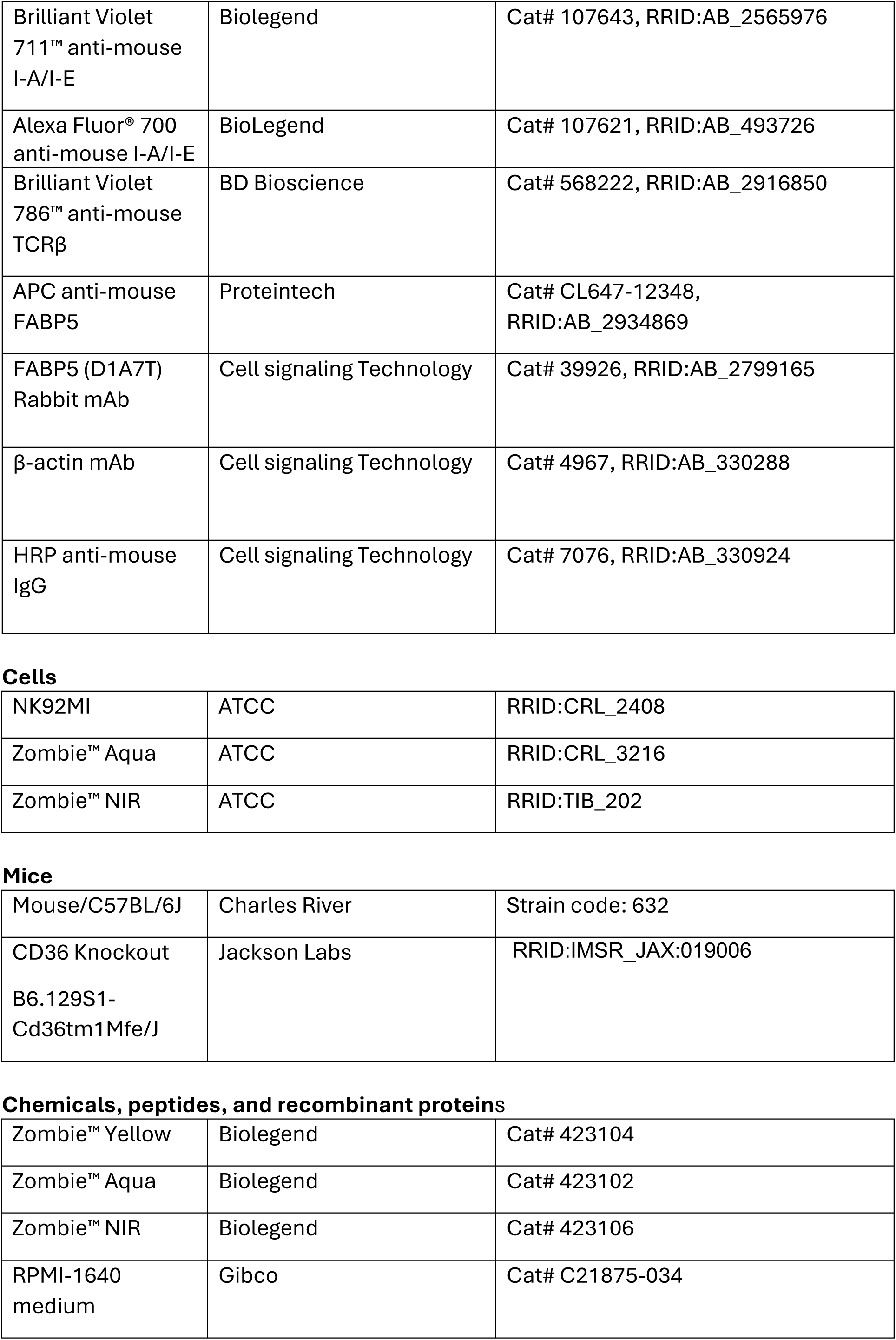
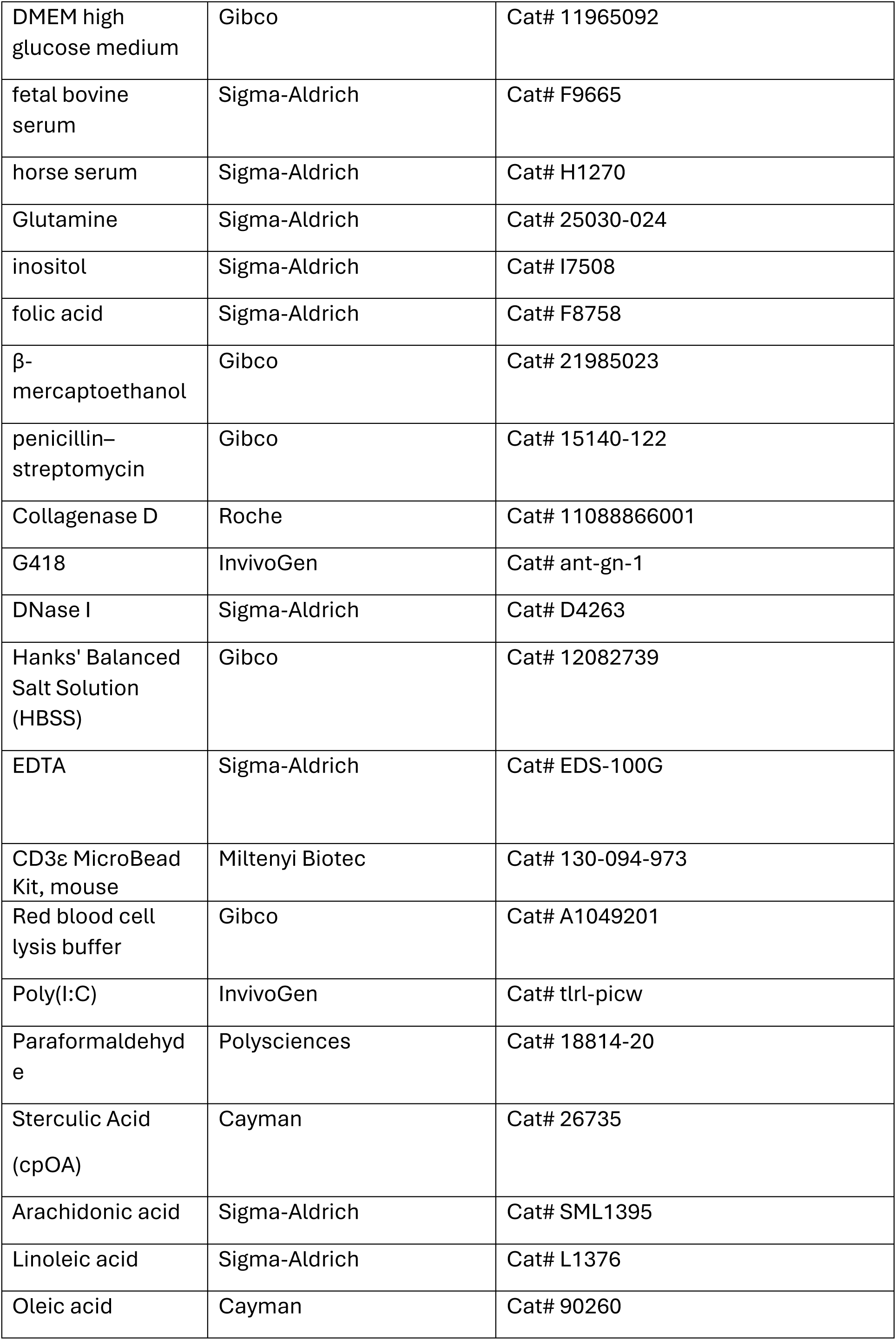
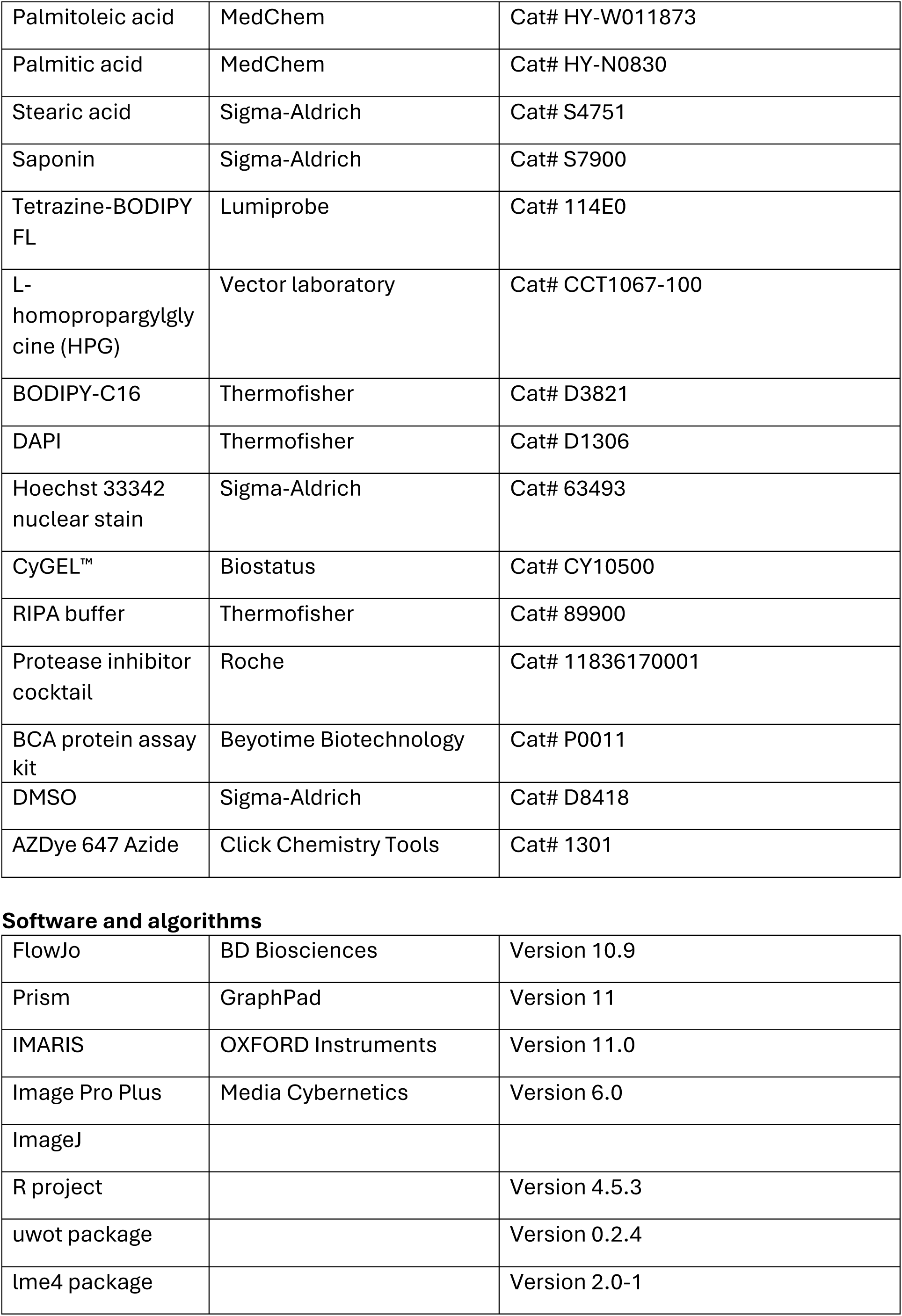
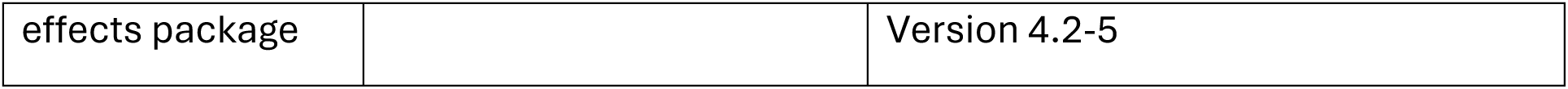

Antibodies
| Antibodies | Source | Cat number |
| --- | --- | --- |
| Fc block | BD Bioscience | Cat# 553141, RRID:AB_394656 |
| Brilliant Violet 510™ anti-mouse CD3 | Biolegend | Cat# 100234, RRID:AB_2562555 |
| Brilliant Violet 750™ anti-mouse CD3 | Biolegend | Cat# 100249, RRID:AB_2734148 |
| Alexa Fluor® 647 anti-mouse CD4 | Biolegend | Cat# 100424, RRID:AB_389324 |
| Brilliant Violet 421™ anti-mouse CD4 | Biolegend | Cat# 100443, RRID:AB_2562557 |
| PE-Cy5 anti-mouse CD8a | BioLegend | Cat# 100709, RRID:AB_312748 |
| APC-Cy7 anti-mouse CD8a | Biolegend | Cat# 100713, RRID:AB_312752 |
| PerCP anti-mouse CD8a | Biolegend | Cat# 100731, RRID:AB_893427 |
| Brilliant Violet 605™ anti-mouse/human CD11b | Biolegend | Cat# 101257, RRID:AB_2565431 |
| PE anti-mouse/human CD11b | Biolegend | Cat# 101207, RRID:AB_312790 |
| PE-Cy7 anti-mouse/human CD11b | Biolegend | Cat# 101215, RRID:AB_312798 |
| PE anti-mouse/human CD11c | BD Bioscience | Cat# 553802, RRID:AB_395061 |
| Brilliant Violet 421™ anti-mouse CD11c | BioLegend | Cat# 117329, RRID:AB_10897814 |
| PE-Cy7 anti-mouse CD19 | BD Bioscience | Cat# 552854, RRID:AB_394495 |
| PE anti-mouse CD19 | Biolegend | Cat# 115507, RRID:AB_313642 |
| Brilliant Violet 650™ CD19 | BD Bioscience | Cat# 563235, RRID:AB_2738085 |
| Super Bright™ 600 anti-mouse CD24 | Thermo Fisher Scientific | Cat# 63-0242-82, RRID:AB_2717023 |
| PerCP-Cy5.5 anti-mouse CD25 | BD Bioscience | Cat# 551071, RRID:AB_394031 |
| PE anti-mouse CD36 | BD Bioscience | Cat# 562702, RRID:AB_2737732 |
| PE-Cy7 anti-mouse CD44 | Thermo Fisher Scientific | Cat# 25-0441-82, RRID:AB_469623 |
| PE anti-mouse CD45 | Biolegend | Cat# 103106, RRID:AB_312971 |
| Brilliant Violet 785™ anti-mouse CD45 | Biolegend | Cat# 103149, RRID:AB_2564590 |
| BD OptiBuild™ BUV805 anti-mouse CD45 | BD Bioscience | Cat# 748370, RRID:AB_2872789 |
| PerCP-Cy5.5 anti-mouse CD45 | BioLegend | Cat# 103132, RRID:AB_893340 |
| APC-Fire™ 810 anti-mouse CD45 | BioLegend | Cat# 103173, RRID:AB_2860599 |
| PE/Dazzle™ 594 anti-mouse CD64 (FcγRI) | BioLegend | Cat# 139319, RRID:AB_2566558 |
| Super Bright™ 702 anti-mouse CD90.2 (Thy-1.2) | Thermo Fisher Scientific | Cat# 67-0902-82, RRID:AB_2717157 |
| APC-eFluor™ 780 anti-mouse CD90.2 (Thy-1.2) | Thermo Fisher Scientific | Cat# 47-0902-82, RRID:AB_1272187 |
| Brilliant Violet 650™ anti-mouse CD98 | BD Bioscience | BD Biosciences Cat# 752888, RRID:AB_2917843 |
| Alexa Fluor® 594 anti-mouse CD172a (SIRPα) | Biolegend | Cat# 144020, RRID:AB_2629588 |
| BD OptiBuild™ BUV805 anti-mouse CD172a (SIRPα) | BD Bioscience | Cat# 741997, RRID:AB_2871296 |
| Brilliant Violet 421™ anti-mouse NK-1.1 | Biolegend | Cat# 108732, RRID:AB_2562218 |
| PE anti-mouse NK-1.1 | Biolegend | Cat# 108707, RRID:AB_313394 |
| PerCP-Cy5.5 anti-mouse NK-1.1 | BioLegend | Cat# 156526, RRID:AB_2894655 |
| PE anti-mouse TCRγδ | Biolegend | Cat# 118107, RRID:AB_313831 |
| PE-Cy7 anti-mouse F4/80 | Biolegend | Cat# 123114, RRID:AB_893478 |
| Brilliant Violet 785™ anti-mouse F4/80 | Biolegend | Cat# 123147, RRID:AB_2564588 |
| PE anti-mouse Ly6C/Ly6G | Thermo Fisher Scientific | Cat# 12-5931-82, RRID:AB_466045 |
| Brilliant Violet 510™ anti-mouse Ly6C | BioLegend | Cat# 128033, RRID:AB_2562351 |
| Spark Blue™ 550 anti-mouse Ly-6G | BioLegend | Cat# 127663, RRID:AB_2860670 |
| Brilliant Violet 785™ anti-mouse/rat XCR1 | Biolegend | Cat# 148225, RRID:AB_2783119 |
| PE-Cy5.5 anti-mouse/rat XCR1 | BioLegend | Cat# 148207, RRID:AB_2564363 |
| Brilliant Violet 711™ anti-mouse I-A/I-E | Biolegend | Cat# 107643, RRID:AB_2565976 |
| Alexa Fluor® 700 anti-mouse I-A/I-E | BioLegend | Cat# 107621, RRID:AB_493726 |
| Brilliant Violet 786™ anti-mouse TCRβ | BD Bioscience | Cat# 568222, RRID:AB_2916850 |
| APC anti-mouse FABP5 | Proteintech | Cat# CL647-12348, RRID:AB_2934869 |
| FABP5 (D1A7T) Rabbit mAb | Cell signaling Technology | Cat# 39926, RRID:AB_2799165 |
| β-actin mAb | Cell signaling Technology | Cat# 4967, RRID:AB_330288 |
| HRP anti-mouse IgG | Cell signaling Technology | Cat# 7076, RRID:AB_330924 |
| Cells |  |  |
| NK92MI | ATCC | RRID:CRL_2408 |
| Zombie™ Aqua | ATCC | RRID:CRL_3216 |
| Zombie™ NIR | ATCC | RRID:TIB_202 |
| Mice |  |  |
| Mouse/C57BL/6J | Charles River | Strain code: 632 |
| CD36 Knockout<br>B6.129S1-Cd36tm1Mfe/J | Jackson Labs | RRID:IMSR_JAX:019006 |
| Chemicals, peptides, and recombinant proteins |  |  |
| Zombie™ Yellow | Biolegend | Cat# 423104 |
| Zombie™ Aqua | Biolegend | Cat# 423102 |
| Zombie™ NIR | Biolegend | Cat# 423106 |
| RPMI-1640 medium | Gibco | Cat# C21875-034 |
| DMEM high glucose medium | Gibco | Cat# 11965092 |
| fetal bovine serum | Sigma-Aldrich | Cat# F9665 |
| horse serum | Sigma-Aldrich | Cat# H1270 |
| Glutamine | Sigma-Aldrich | Cat# 25030-024 |
| inositol | Sigma-Aldrich | Cat# I7508 |
| folic acid | Sigma-Aldrich | Cat# F8758 |
| β-mercaptoethanol | Gibco | Cat# 21985023 |
| penicillin–streptomycin | Gibco | Cat# 15140-122 |
| Collagenase D | Roche | Cat# 11088866001 |
| G418 | InvivoGen | Cat# ant-gn-1 |
| DNase I | Sigma-Aldrich | Cat# D4263 |
| Hanks' Balanced Salt Solution (HBSS) | Gibco | Cat# 12082739 |
| EDTA | Sigma-Aldrich | Cat# EDS-100G |
| CD3ε MicroBead Kit, mouse | Miltenyi Biotec | Cat# 130-094-973 |
| Red blood cell lysis buffer | Gibco | Cat# A1049201 |
| Poly(I:C) | InvivoGen | Cat# tlrl-picw |
| Paraformaldehyde | Polysciences | Cat# 18814-20 |
| Sterculic Acid (cpOA) | Cayman | Cat# 26735 |
| Arachidonic acid | Sigma-Aldrich | Cat# SML1395 |
| Linoleic acid | Sigma-Aldrich | Cat# L1376 |
| Oleic acid | Cayman | Cat# 90260 |
| Palmitoleic acid | MedChem | Cat# HY-W011873 |
| Palmitic acid | MedChem | Cat# HY-N0830 |
| Stearic acid | Sigma-Aldrich | Cat# S4751 |
| Saponin | Sigma-Aldrich | Cat# S7900 |
| Tetrazine-BODIPY FL | Lumiprobe | Cat# 114E0 |
| L-homopropargylglycine (HPG) | Vector laboratory | Cat# CCT1067-100 |
| BODIPY-C16 | Thermofisher | Cat# D3821 |
| DAPI | Thermofisher | Cat# D1306 |
| Hoechst 33342 nuclear stain | Sigma-Aldrich | Cat# 63493 |
| CyGEL™ | Biostatus | Cat# CY10500 |
| RIPA buffer | Thermofisher | Cat# 89900 |
| Protease inhibitor cocktail | Roche | Cat# 11836170001 |
| BCA protein assay kit | Beyotime Biotechnology | Cat# P0011 |
| DMSO | Sigma-Aldrich | Cat# D8418 |
| AZDye 647 Azide | Click Chemistry Tools | Cat# 1301 |
| Software and algorithms |  |  |
| FlowJo | BD Biosciences | Version 10.9 |
| Prism | GraphPad | Version 11 |
| IMARIS | OXFORD Instruments | Version 11.0 |
| Image Pro Plus | Media Cybernetics | Version 6.0 |
| ImageJ |  |  |
| R project |  | Version 4.5.3 |
| uwot package |  | Version 0.2.4 |
| lme4 package |  | Version 2.0-1 |
| effects package |  | Version 4.2-5 |

---

**Supplementary Figure 1.**
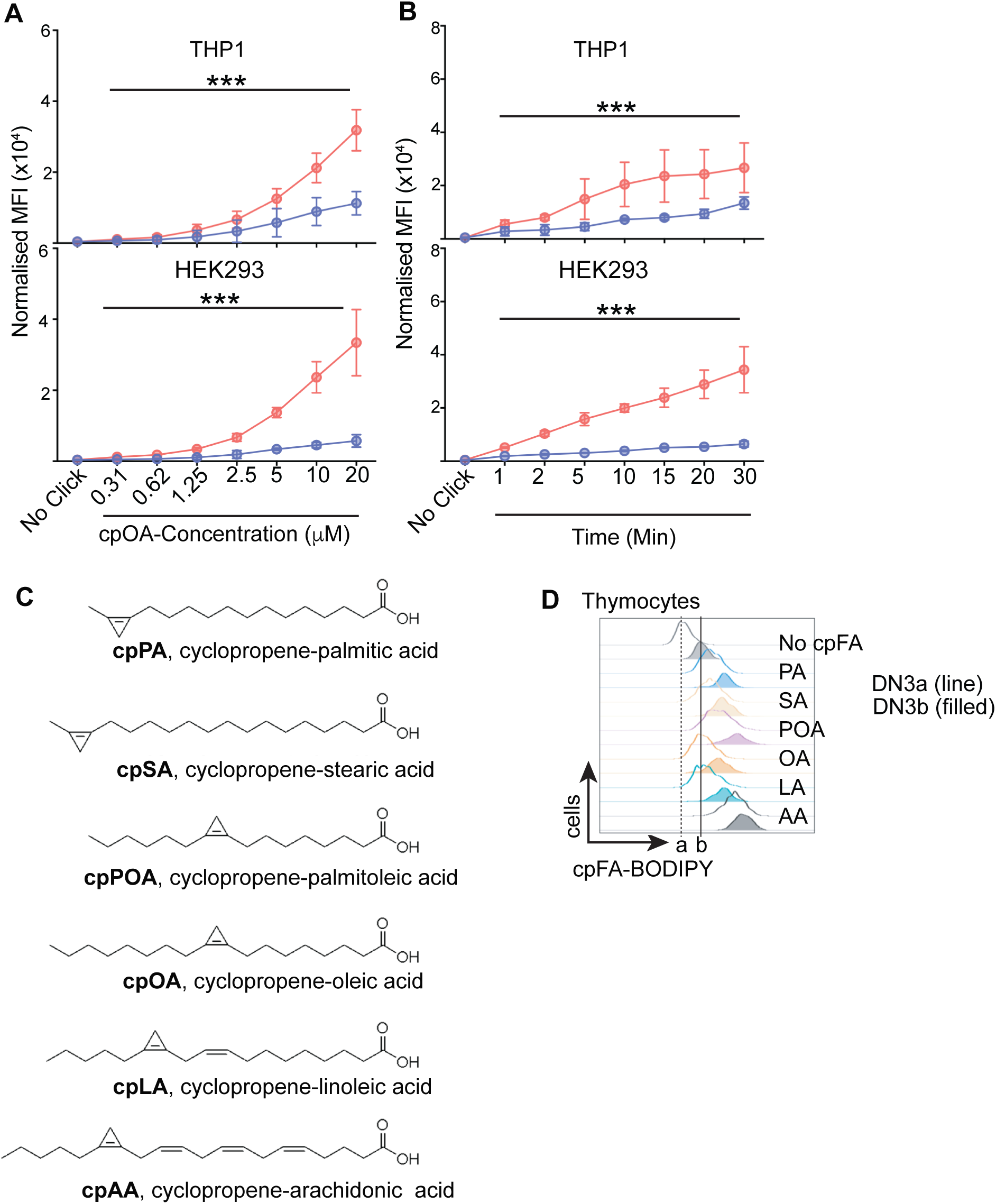
(**A,B)** cpOA uptake assays were performed in THP-1(top) and HEK293 (bottom) cells for 10 min using increasing concentrations of cpOA, as indicated (A), and using (10 μM) cpOA and increasing incubation times, as indicated (B). **(C)** Structures of cpFA molecules used in this study. **(D)** Thymocytes were isolated and uptake of 6 different cpFAs (10 μM for 10 min) measured in DN3a (outlined histograms) and DN3b (filled histograms) thymocytes. Representative histograms of uptakes are shown. Dotted and filled lines indicate no cpFA background controls for DN3a and DN3b subsets, respectively. Data is representative (D) or mean +/- SEM (A,B) for 4 (D) mice and 3 (B), 4 (B,D) independent experiments. Statistical analysis was performed using ANOVA. ***, p < 0.001; SA, steric acid; PA, palmitic acid; OA, oleic acid; POA, palmitoleic acid; LA, linoleic acid; AA, arachidonic acid.

**Supplementary Figure 2.**
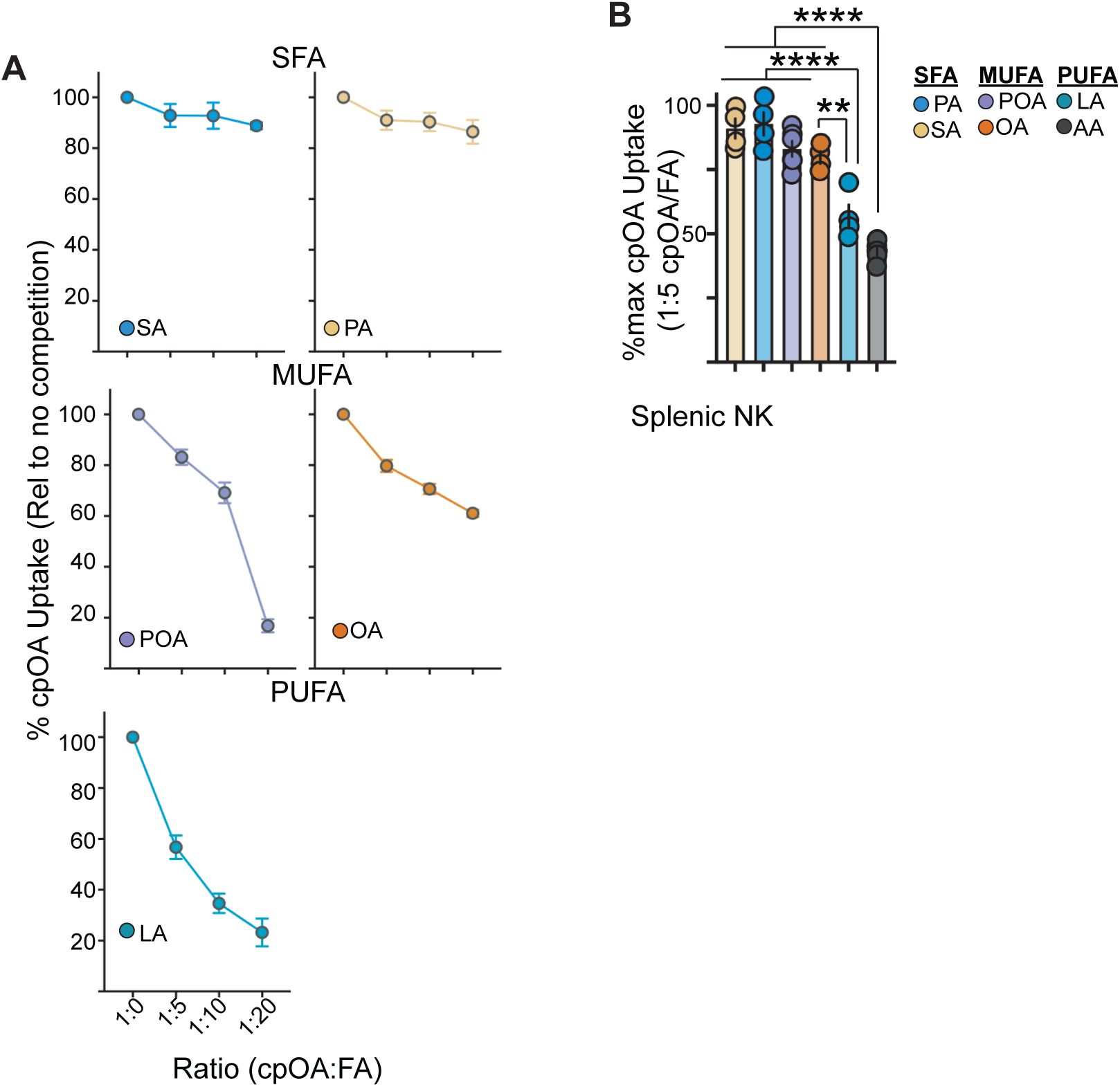
**(A,B)** Splenocytes were incubated with cpOA (10 μM) for 10 min at 37°C in the presence of varying concentrations of native OA or native forms of other FA. Pooled data for cpOA uptake into splenic NK cells with increasing ratios of different FA species (A). Pooled data for the percentage of max cpOA uptake observed with a 1:5 (cpOA:FA) ratio of each native FA species. Data is mean +/- SEM for 4-6 mice from at least 3 independent experiments. Statistical analysis was performed using ANOVA and Tukey (B) post-tests. **, p < 0.01; ****, p < 0.0001; SFA, saturated FA; MUFA, mono-unsaturated FA; PUFA, poly-unsaturated FA; SA, steric acid; PA, palmitic acid; OA, oleic acid; POA, palmitoleic acid; LA, linoleic acid; AA, arachidonic acid.

**Supplementary Figure 3.**
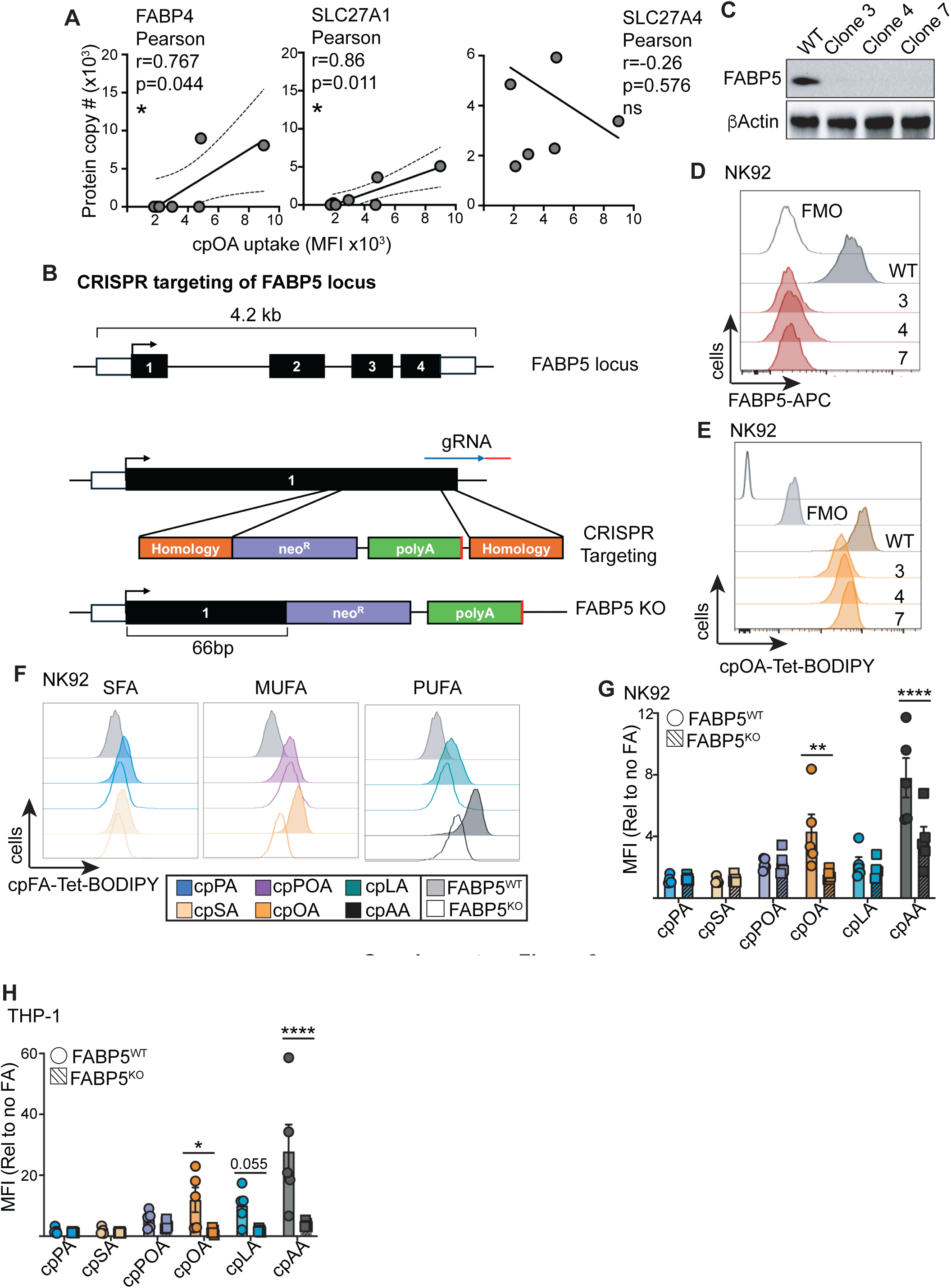
**(A)** Protein copy numbers for proteins associated with FA transport for murine splenocyte populations and lymph node T cells from proteomic datasets (immpres.co.uk, PRIDE) for FABP4, SLC27A1 and SLC27A4 graphed with corresponding cpOA uptake measured by flow cytometry. **(B)** Outline of CRISPR targeting strategy for deleting FABP5 from NK92, THP-1 cell lines. **(C)** Representative western blot of WT NK92 cells and 3 separate clones of FABP5 KO NK92 cells. **(D,E)** Flow cytometry analysis of FAPB5 expression (D) and cpOA uptake (E) in WT NK92 cells and 3 separate clones of FABP5 KO NK92 cells. **(F,G,H)** Flow cytometry analysis of the uptake of 6 different cpFA probes (10 μM for 10 min) into NK92 cells (F,G) and THP-1 cells (H) comparing FABP5 WT and KO cells. Data is representative (C-F), mean (A), mean +/- SEM (G,H) for 3-4 proteomic and uptake replicates (A), and 3 (C-E), 5 (G) and 3 (H) independent experiments. Statistical analysis was performed using ANOVA and Tukey post-tests or using a Pearson’s correlation test (A). ns, not significant; *, p < 0.05; **, p < 0.01; ****, p < 0.0001. SA, steric acid; PA, palmitic acid; OA, oleic acid; POA, palmitoleic acid; LA, linoleic acid; AA, arachidonic acid.

**Supplementary Figure 4.**
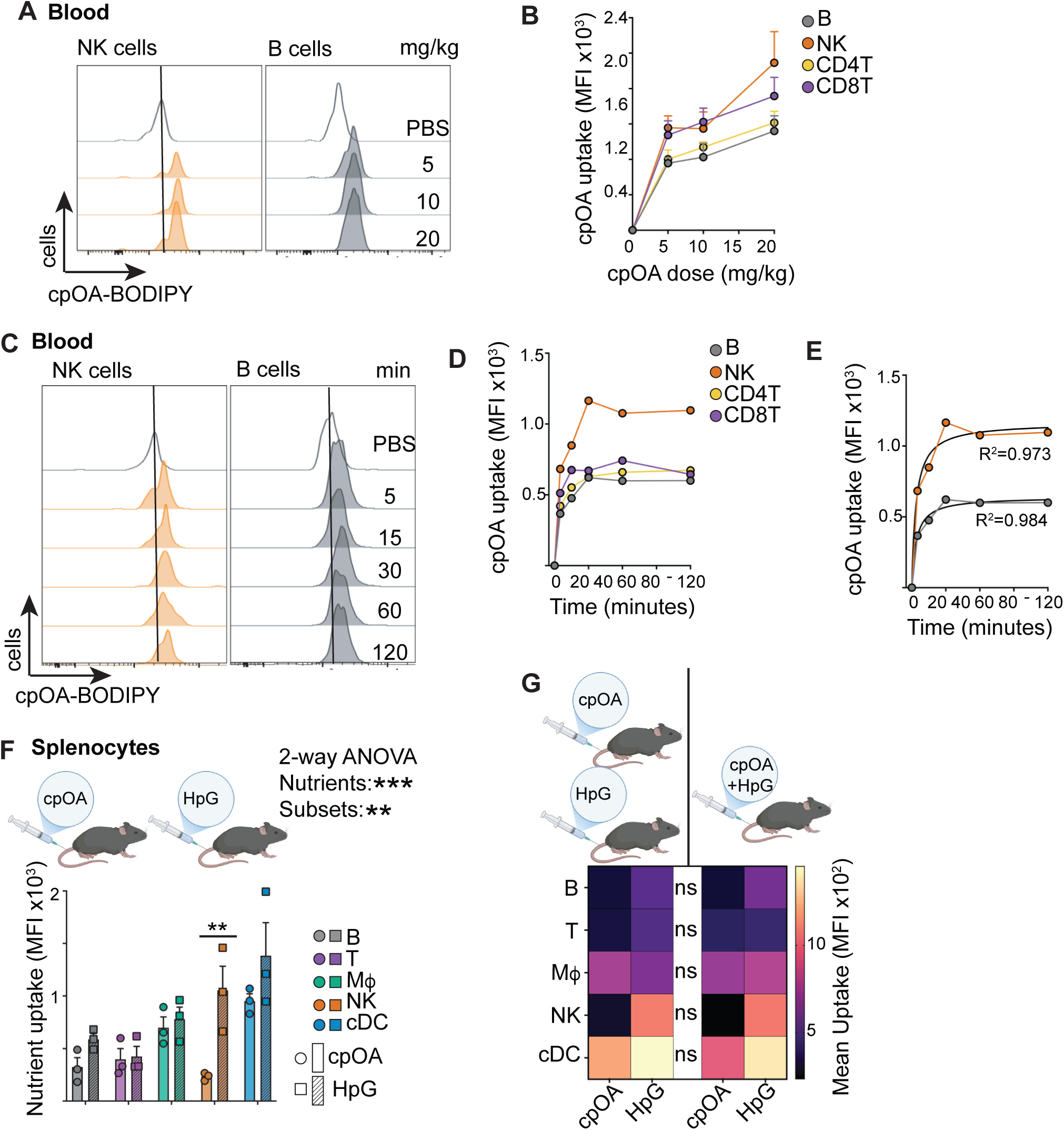
**(A-E)** Mice were i.v. injected with cpOA at indicated doses (A,B) and with 10 mg/kg for indicated times (C,D) or PBS controls, and blood isolated, cells prepared for flow cytometry analysis. Representative histograms are shown for NK cells and B cells (A,C) and pooled data for CD4^+^ T cells, CD8^+^ T cells, NK cells and B cells (B,D). Uptake kinetics for NK cells and B cells were modelled to hyperbolic function and R^2^ values shown (E). (**F,G)** Mice were i.v. injected with 10 mg/kg cpOA, 100 mg/kg HPG,10 mg/kg cpOA and 100 mg/kg HPG together, or PBS control; then after 30 min spleens were isolated and cells prepared for flow cytometry analysis. **(F)** Pooled data showing cpOA and HpG uptake into splenic subsets in separate mice after single click-probe injections and single click reactions prior to flow cytometry analysis. **(G)** Heatmap comparing the mean uptake values for cpOA and HpG uptake into splenic immune cells when cpOA and HpG were injected into separate mice with single click reactions performed, or together into one mouse, with double click reactions performed prior to analysis. Data is representative (A,C) and mean +/- SEM (B,D,E,F) and mean (G) for 3 (A-G) separate mice and 3 individual experiments. Statistical analysis was performed using ANOVA or non-parametric tests as appropriate; Šidák (F) and Tukey (G) post-tests or analysed using non-linear regression to hyperbola (E). ns, not significant; **, p < 0.01; ***, p < 0.001.

